# Psychedelic-mediated Reversal of General Anesthesia and Restoration of Brain Dynamics in Rat

**DOI:** 10.1101/2025.01.22.632279

**Authors:** Emma R. Huels, Nicholas Kolbman, Christopher W. Fields, Amanda G. Nelson, Hazel Jackson, Tiecheng Liu, George A. Mashour, Dinesh Pal

## Abstract

Serotonergic psychedelics enhance neurophysiological complexity and the repertoire of brain states, whereas general anesthetics produce opposite effects. Serotonergic psychedelics are also known to increase wakefulness and reduce sleep time in rodents. Therefore, we hypothesized that 2,5-dimethoxy-4-iodopamphetamine (DOI), a serotonergic psychedelic, will reverse general anesthesia and restore neurophysiological conditions associated with normal wakefulness. We demonstrate that intravenous administration of DOI in rats under general anesthesia induced wakefulness despite ongoing delivery of the anesthetics, propofol or isoflurane. Behavioral arousal was accompanied by recovery of directional and non-directional high gamma (125-165Hz) functional connectivity and restoration of functional brain network structure. These behavioral and neurophysiological effects were blocked by a 5-HT2A antagonist, volinanserin. Intravenous administration of a non-psychedelic 5-HT2A agonist, lisuride, failed to restore wakefulness or brain dynamics in anesthetized rats. To our knowledge, these results provide the first evidence of psychedelic-mediated reversal of general anesthesia and concurrent restoration of brain dynamics associated with normal wakefulness.

## Main Text

Despite decades of previous work—including studies that have explored psychostimulants such as nicotine, methylphenidate, amphetamine, and caffeine to reverse or accelerate recovery from the anesthetized state^1–7^—there is currently no standard reversal agent for general anesthesia or sedation in the clinical setting. Furthermore, previous studies focused primarily on behavioral assessment and did not investigate functional brain dynamics often observed during normal wakefulness, which can provide a translational bridge to human studies, including anesthetized patients and those with disorders of consciousness.

Serotonergic psychedelics such as 2,5-dimethoxy-4-iodoamphetamine (DOI) act primarily through excitatory 5-HT2A receptors,^8^ which are highly concentrated within the medial prefrontal cortex^9–14^—a region previously shown to play a role in mediating arousal,^15–17^ including reversal of general anesthesia.^18^ Additionally, accumulating evidence suggests that general anesthesia and psychedelics induce opposite effects at the global brain network level. General anesthetics decrease neurophysiological complexity and contract the repertoire of functional brain states^19–22^ whereas psychedelics increase neurophysiological complexity and expand the repertoire of functional brain states.^23,24^ A potentially arousal-promoting effect of serotonergic psychedelics is also supported by studies reporting an acute increase in wakefulness and suppression of sleep after systemic administration.^25–28^ Based on these data, we hypothesized that systemic administration of DOI will reverse general anesthesia in rats and restore functional brain dynamics—measured via high-density electroencephalogram (EEG) recordings—associated with normal wakefulness. In addition, we used pretreatment with a 5-HT2A antagonist, volinanserin, to determine whether DOI acts via 5-HT2A receptors to reverse anesthesia and restore brain dynamics. Finally, we conducted additional studies to determine whether a non-psychedelic 5-HT2A agonist, lisuride, can mimic the effect of the psychedelic 5-HT2A agonist DOI on behavioral arousal and restoration of brain dynamics.

## Results

All studies were conducted using a within-subject approach designed to allow each rat to participate in all three experimental sessions, i.e., saline infusion during anesthesia, DOI infusion during anesthesia, and DOI infusion preceded by pretreatment with volinanserin during anesthesia. The same within-subject approach was used for a second cohort of rats undergoing saline infusion during anesthesia, DOI infusion during anesthesia, and lisuride infusion during anesthesia. To avoid any potential bias arising from following a particular order of experimental sessions, the experimental sessions were conducted in a random manner.

### Intravenous Administration of DOI Restored Wakefulness during General Anesthesia

A schematic illustrating the experimental timeline is provided in **Fig. 1A**. The onset of behavioral arousal after intravenous DOI administration during propofol or isoflurane anesthesia (i.e., active emergence) typically began within one minute after cessation of the DOI infusion. Arousal scores, assessed by a reviewer blinded to the experimental conditions (see *Arousal Scoring Criteria* in Materials and Methods), were significantly higher after DOI administration, as compared to saline infusion, during propofol or isoflurane anesthesia (*p* < .0001) (**Fig. 1B**). Pretreatment with volinanserin completely abolished this effect, resulting in behavioral scores that were indistinguishable from those observed after saline administration (*p* = 1.00), and decreased the arousal score as compared to DOI only (i.e., without volinanserin pretreatment) (*p* < .0001) (**Fig. 1B**). Representative behavior showing the return of righting reflex (RORR, arousal score=4) following infusion of DOI during propofol or isoflurane anesthesia is provided in **movie S1** and **movie S2**, respectively.

**Fig. 1.**
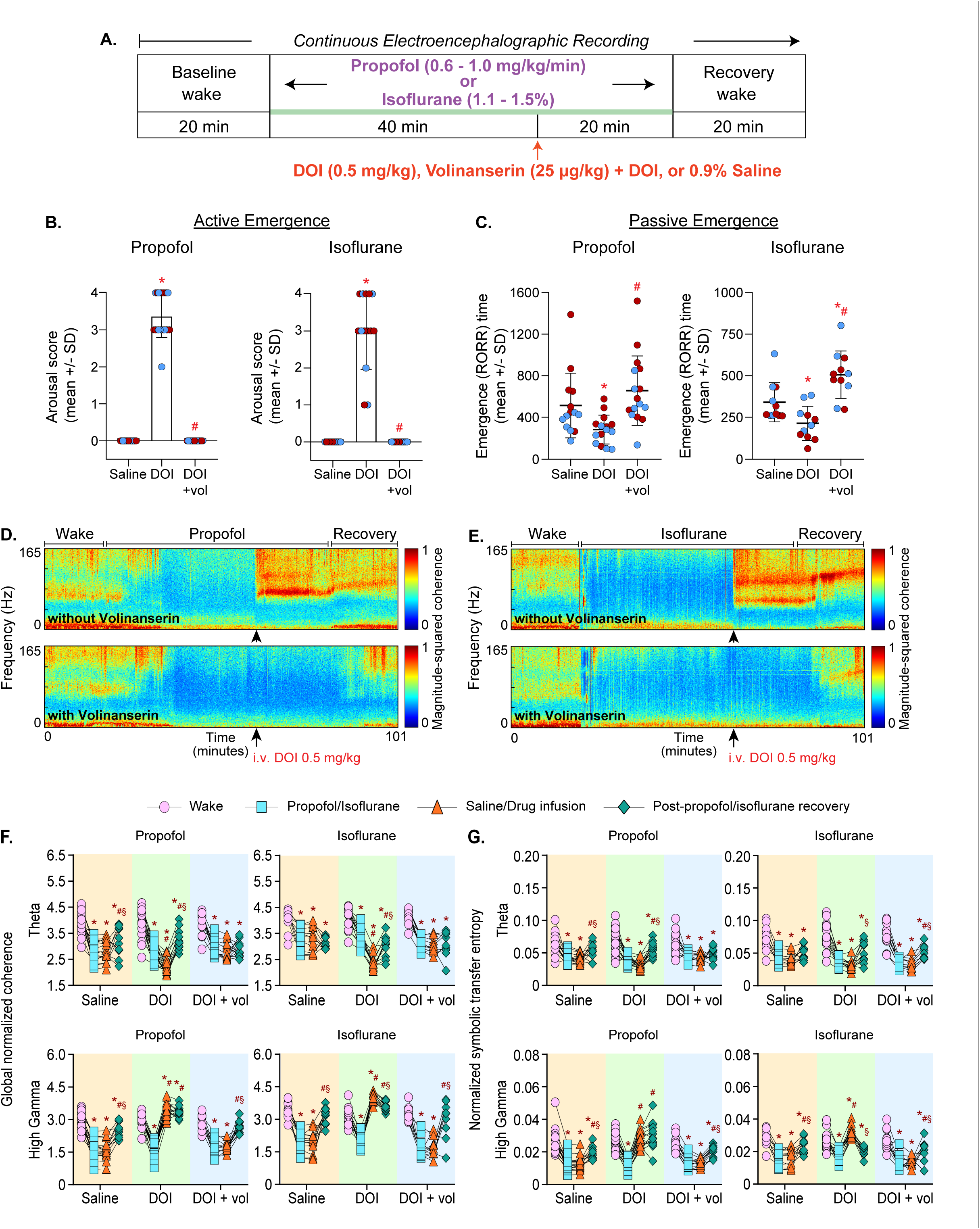
Intravenous Administration of 2,5-Dimethoxy-4-Iodopamphetamine (DOI) Reversed General Anesthesia and Restored Functional Connectivity via 5-HT2A Receptors. **A.** Schematic showing experiment timeline. All experiments were conducted using a within subject design. **B.** Intravenous administration of DOI induced active reversal of general anesthesia as demonstrated by the increase in the arousal score during propofol or isoflurane anesthesia whereas intravenous pretreatment with volinanserin blocked the arousal promoting effect of DOI. Arousal Scores: 1 – EEG activation, 2 – coordinated movements, 3 – attempts to regain righting reflex, 4 – return of righting reflex (RORR) C. DOI facilitated passive emergence as evident from a decrease in the time to return of righting reflex (RORR) after cessation of propofol or isoflurane anesthesia. Intravenous pretreatment with volinanserin blocked the effect of DOI in accelerating passive emergence. For **B-C**, significance symbols (*p* < .05) show post hoc pairwise tests between conditions that were performed with single-step correction for multiple comparisons via Tukey’s test: *compared to saline infusion, #compared to DOI infusion. Exact *p* values are reported in the results section and supplementary tables S1-S2. **D-E**. Averaged cohereograms illustrating the global coherence across frequencies (0.1-165 Hz) before, during, and after propofol (**D**) or isoflurane (**E**) anesthesia during which the rats received infusion of DOI or DOI with volinanserin pretreatment. The black arrow indicates the time of DOI delivery. Warmer colors indicate higher coherence while cooler colors indicate less coherence. **F-G**. Line-symbol plots show global normalized coherence (**F**) and feedback connectivity (measured via normalized symbolic transfer entropy, **G**) for each rat within the theta (4-10 Hz) and high gamma (125-165 Hz) band during wake, anesthesia (propofol or isoflurane), post-drug administration (saline, DOI, or DOI with volinanserin pretreatment during propofol or isoflurane anesthesia), and post-anesthetic recovery after the return of righting reflex. Each data point (pink circle – wake; blue square – anesthesia; orange triangle – anesthesia + saline, DOI, or DOI with volinanserin pretreatment; green diamond – post-anesthetic recovery) shows the global normalized coherence (**F**) or feedback connectivity (**G**) for an individual rat. Significance symbols (*p* < .05) show post hoc pairwise tests between states that were performed with single-step correction for multiple comparisons via Tukey’s test: *compared to wake, #compared to anesthesia only, §compared to saline or drug (DOI or DOI pretreated with volinanserin) administration during anesthesia. Exact *p* values are reported in the results section and supplementary tables S7-10 and S15-18.

While our primary objective was to assess the effect of the drug/saline infusion on active reversal of general anesthesia, we also assessed passive emergence from general anesthesia (i.e., the time to RORR after the anesthetic was discontinued) (**Fig. 1C**). Therefore, rats which previously achieved RORR after the drug infusion (i.e., active reversal) were returned to a supine position during the last 5 minutes of anesthetic delivery and the time to RORR following anesthetic cessation was measured. As compared to saline infusion, infusion of DOI decreased the time to RORR (i.e., passive emergence) after cessation of propofol (*p* = .0088) or isoflurane (*p* = .042) anesthesia. Similar to the effect on active emergence, pretreatment with volinanserin blocked this effect and resulted in righting times that were similar to the saline condition in the propofol group (*p* = .43), and increased righting times as compared to saline infusion in the isoflurane group (*p* = .0071). Passive emergence times following pretreatment with volinanserin were also increased compared to DOI alone in the propofol (*p* = .00023) and isoflurane (*p* < .0001) groups (**Fig. 1C**).

Across experimental groups (i.e., saline, DOI, DOI pretreated with volinanserin), there was no statistical difference in arousal scores between female and male rats during propofol or isoflurane anesthesia (*p* < 1.00). However, female rats took longer than males to regain the righting reflex after the end of propofol anesthesia (*p* = .015); there was no statistical difference in the time to return of righting reflex between female and male rats after isoflurane anesthesia (*p* = .069).

### Intravenous Administration of DOI Restored High Gamma, but not Theta, Non-Directional Connectivity

Based on previous reports from our and other laboratories showing an association between arousal states and activity in theta (4-10 Hz) and high gamma (125-165 Hz) bands,^29–32^ we focused our EEG analyses on these two frequency bands. Data on the power spectral changes are provided in **Fig. S1A-C** and the supplementary section. Continuous coherograms illustrating changes in global coherence before and after DOI administration during propofol or isoflurane anesthesia with or without volinanserin pretreatment are provided in **Fig. 1D-E**. As expected, theta global coherence decreased during propofol or isoflurane anesthesia (*p* < .0001) (**Fig. 1F**). Unexpectedly, theta global coherence decreased further after the infusion of DOI during propofol or isoflurane anesthesia (*p* < .0001). As compared to theta global coherence during anesthesia, there was no change in global coherence after saline infusion (*p* ≤ 1.00) or when DOI was infused after volinanserin pretreatment (*p* < 1.00). Theta global coherence remained low (compared to wake) during the recovery period across all three conditions (i.e., saline infusion, infusion of DOI, and infusion of volinanserin followed by DOI infusion) for propofol and isoflurane groups (*p* < .0001) (**Fig. 1F**).

High gamma global coherence decreased during propofol or isoflurane anesthesia (*p* < .0001) (**Fig. 1F**). DOI infusion reversed these changes in high gamma global coherence during anesthesia induced by either anesthetic, such that the global coherence values exceeded waking levels (DOI vs. wake, *p* < .0001). Pretreatment with volinanserin blocked the DOI-induced changes in high gamma global coherence during propofol or isoflurane anesthesia (*p* < 1.00), with similar results occurring after the saline infusion for each anesthetic group (*p* ≤ 1.00). High gamma global coherence returned to waking levels during the recovery period for all conditions in the isoflurane group (*p* < 1.00). Following the cessation of propofol delivery and RORR, the high gamma global coherence remained elevated (compared to wake) in the rats that received DOI (*p* = .01), decreased in the rats that received saline (*p* < .0001), and returned to baseline waking levels in the rats who received DOI following pretreatment with volinanserin (*p* = .16) (**Fig. 1F**).

### Intravenous Administration of DOI Restored High Gamma, but not Theta, Directional Connectivity

Next, we measured changes in feedback (frontal-to-parietal) and feedforward (parietal-to-frontal) normalized symbolic transfer entropy (NSTE)—a measure of directed functional connectivity—between frontal and parietal electrode pairs. Theta feedback connectivity decreased during anesthesia induced by either propofol or isoflurane (*p* < .0001) (**Fig. 1G**). As compared to theta feedback connectivity during propofol or isoflurane anesthesia, there was no significant change in theta feedback connectivity following DOI infusion during propofol (*p* = .28) or isoflurane (*p* = .34) anesthesia. There was also no statistical change following the infusion of saline or DOI with volinanserin pretreatment during either anesthetic (*p* ≤ 1.00). Compared to feedback connectivity during baseline wake state, theta feedback connectivity remained decreased during the post-propofol recovery period across conditions (saline: *p* = .051; DOI: *p* = .0015; DOI + volinanserin: *p* = .00029), with similar results during the post-isoflurane recovery period across conditions (*p* < .0001) (**Fig. 1G**).

Across conditions, high gamma feedback connectivity decreased during propofol or isoflurane anesthesia (*p* < .0001) (**Fig. 1G**). DOI infusion during either propofol or isoflurane anesthesia restored high gamma feedback connectivity to or above waking levels (for propofol, DOI vs. wake: *p* = .42; for isoflurane, DOI vs. wake: *p* = .0079). High gamma feedback connectivity remained at waking levels during the post-propofol recovery period (*p* = .15) and decreased slightly during the post-isoflurane recovery period (*p* = .0051) in the rats that received DOI. No statistical change in high gamma feedback connectivity occurred in the rats that received saline infusion (*p* ≤ 1.00) or DOI infusion after pretreatment with volinanserin (*p* = 1.00) during propofol or isoflurane anesthesia. As compared to wake state, high gamma feedback connectivity remained decreased during the post-anesthesia recovery wake period in the rats that received saline infusion or infusion of DOI after volinanserin (for propofol, saline: *p* = .0001; DOI + volinanserin: *p* = .049; for isoflurane, saline: *p* = .044; DOI + volinanserin: *p* = .0089) (**Fig. 1G**). The changes in feedforward connectivity were similar to that reported here for feedback connectivity and are provided in **Figure S1D**.

### Intravenous Administration of DOI Restored High Gamma, but not Theta, Node Degree

To assess network-level changes induced by DOI infusion during general anesthesia, we calculated node degree (i.e., for a given electrode, the number of other electrodes to which it is functionally connected). Topographic heat maps in **Fig. 2A-B** illustrate the average node degree across rats for each electrode within each state, frequency band, and anesthetic for each condition. Theta node degree decreased during propofol (all conditions *p* < .0001) or isoflurane (for saline, *p* = .021; for DOI, *p* = .00061; for DOI + volinanserin, *p* < .0001) anesthesia and decreased further after infusion of DOI during either anesthetic (*p* < .0001); the decrease in node degree after DOI was blocked by pretreatment with volinanserin (*p* < 1.00) and was not observed after the saline infusion (*p* < 1.00). Compared to wakefulness, theta node degree remained decreased across all conditions during the post-propofol (for saline, *p* = .0024; for DOI, *p* = .0018; for DOI + volinanserin, *p* < .0001) and post-isoflurane (all conditions *p* < .0001) recovery period (**Fig. 2C**). High gamma node degree also decreased during propofol or isoflurane anesthesia across conditions (*p* < .0001). In contrast to theta node degree which decreased after DOI infusion, node degree in the high gamma band increased above waking levels after DOI infusion (for propofol, *p* < .0001; for isoflurane, *p* = .015) and remained elevated in the post-anesthesia period after the cessation of propofol but not isoflurane anesthesia (for propofol recovery vs. wake, *p* = .00069; for isoflurane recovery vs. wake, *p* = .070). No statistical change in high gamma node degree occurred after saline infusion (*p* < 1.00) or infusion of DOI following pretreatment with volinanserin (*p* ≤ 1.00). For both control conditions, high gamma node degree returned to waking levels during the post-isoflurane (for saline, *p* < 1.00; for DOI + volinanserin, *p* = .45) but not post-propofol (for saline: *p* < .0001; for DOI + volinanserin: *p* = .035) recovery period (**Fig. 2C**).

**Fig. 2.**
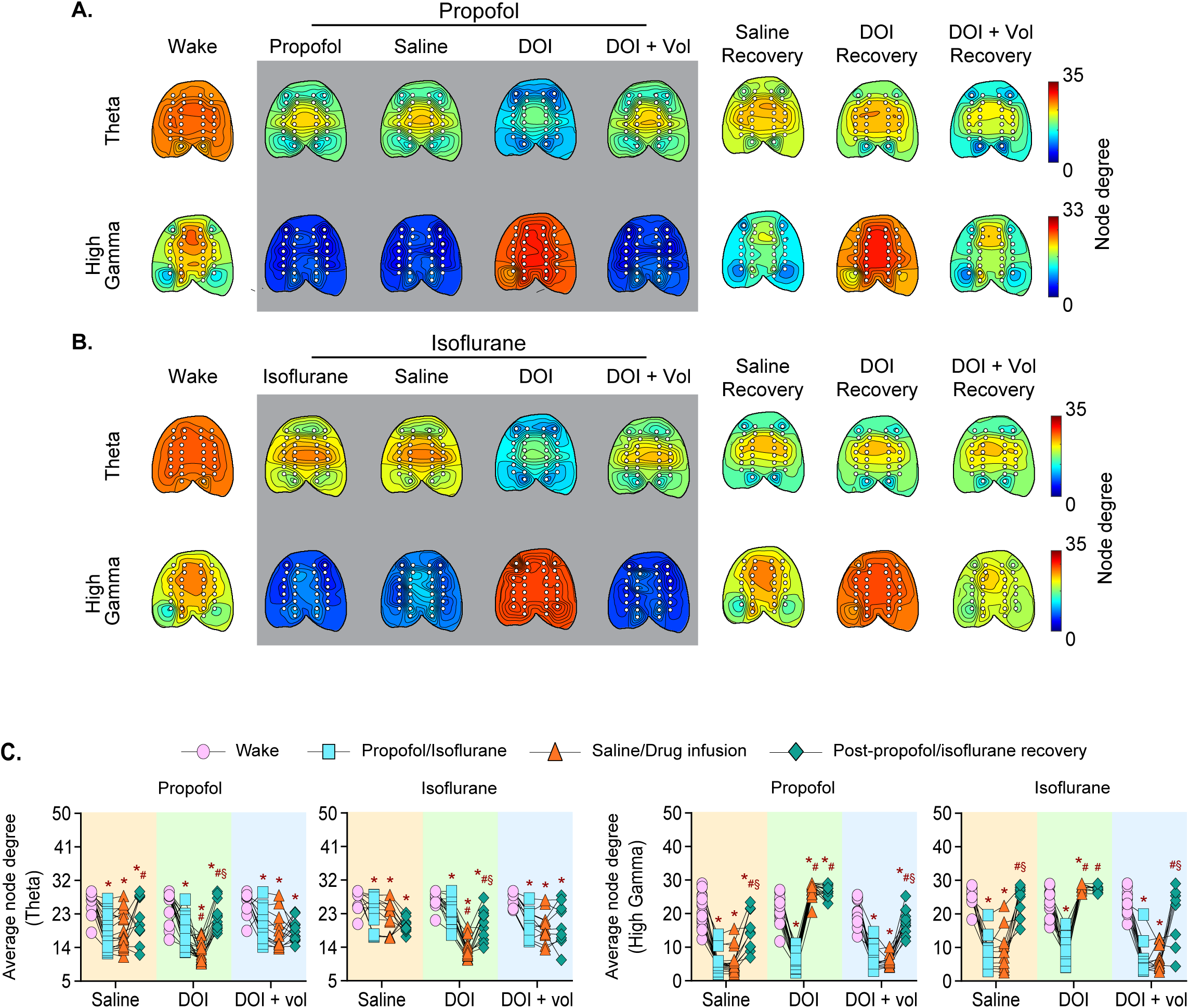
Intravenous Administration of 2,5-Dimethoxy-4-Iodopamphetamine (DOI) Restored High Gamma, but not Theta, Node Degree. **A-B.** Topographic maps averaged across all rats illustrate the node degree for each electrode during wake, anesthesia (propofol or isoflurane), post-drug administration (saline, DOI, or DOI with volinanserin pretreatment during propofol or isoflurane anesthesia), and post-anesthetic recovery after the return of righting reflex. Warmer colors indicate higher node degree whereas cooler colors indicate lower node degree. Each white circle represents a single electrode, with 30 electrodes plotted on each topographic map. **C.** Line-symbol plots showing average node degree across electrodes for each rat within the theta (4-10 Hz) and high gamma (125-165 Hz) bands during wake, anesthesia (propofol or isoflurane), post-drug administration (saline, DOI, or DOI with volinanserin pretreatment during propofol or isoflurane anesthesia), and post-anesthetic recovery after the return of righting reflex. Each data point (pink circle – wake; blue square – anesthesia; orange triangle – anesthesia + saline, DOI, or DOI with volinanserin pretreatment; green diamond – post-anesthetic recovery) shows the average node degree for individual rats. Significance symbols (*p* < .05) show post hoc pairwise tests between states that were performed with single-step correction for multiple comparisons via Tukey**’s** test: *compared to wake, #compared to anesthesia only, §compared to saline or drug (DOI or DOI pretreated with volinanserin) administration during anesthesia. Exact *p* values are reported in the results section and supplementary tables S19-S22.

### Intravenous Administration of Lisuride, a Non-Psychedelic 5-HT2A Agonist, during General Anesthesia Did Not Restore Wakefulness

In a separate cohort of rats, we investigated whether the intravenous infusion of lisuride, a non-psychedelic 5-HT2A partial agonist, could mimic the arousal-promoting effects of DOI (**Fig. 3A**). Since DOI infusion had a similar effect on behavioral arousal during both propofol and isoflurane anesthesia, the experiments to test the effect of lisuride on behavioral arousal were conducted only during propofol anesthesia. As compared to saline infusion, infusion of lisuride did not produce any statistically significant change in the arousal scores (*p* < 1.00). To confirm that the lack of arousal after lisuride infusion is not due to any technical confounds, we administered DOI intravenously in these rats during propofol anesthesia, which did produce behavioral arousal and increased arousal score as compared to saline infusion (*p* < .0001) (**Fig. 3B**).

**Fig. 3.**
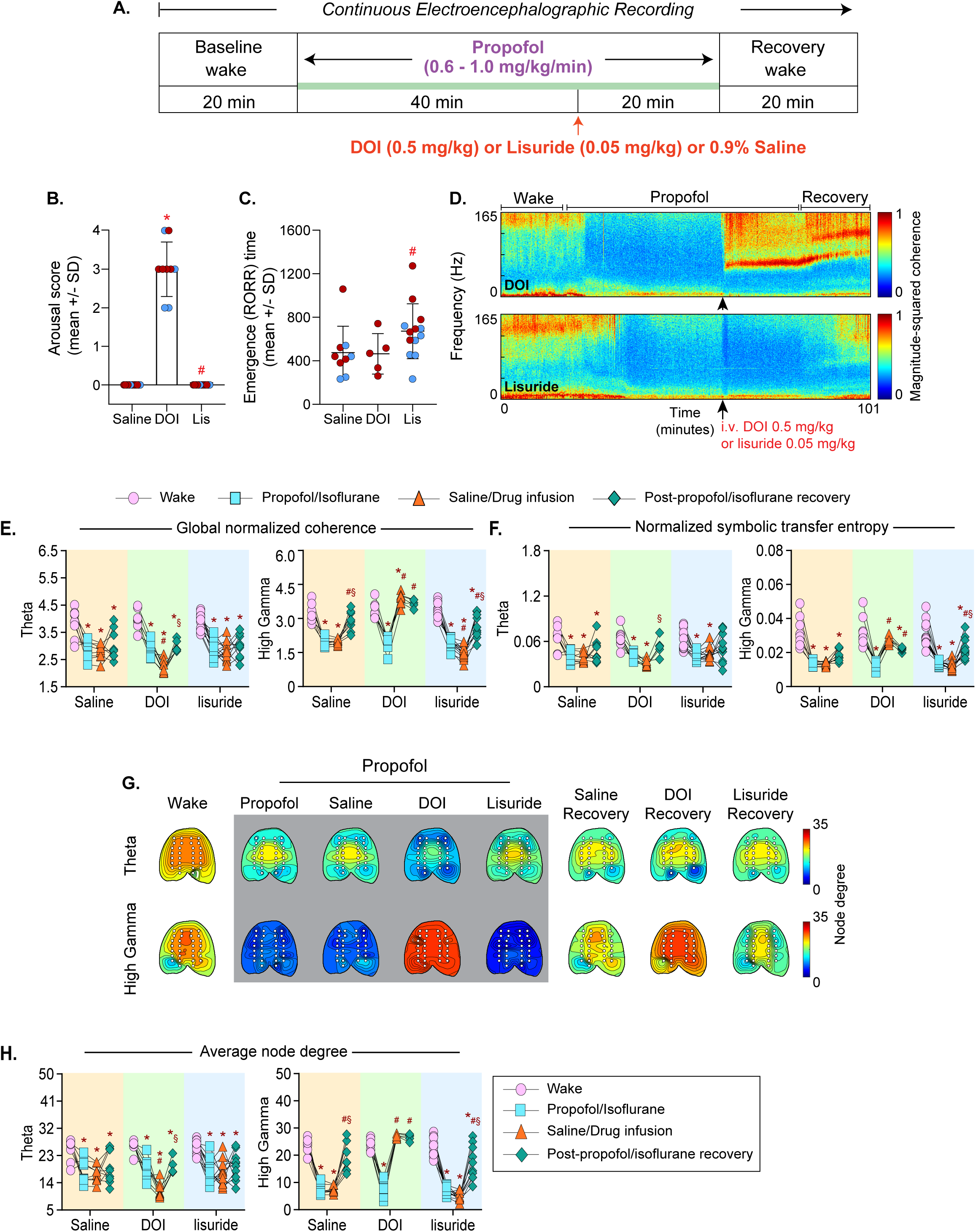
Intravenous Administration of Lisuride Failed to Reverse Propofol Anesthesia and Did Not Restore Functional Connectivity or Brain Network Structure. **A.** Schematic showing experiment timeline. All experiments were conducted using a within subject design. **B.** Unlike DOI, intravenous administration of lisuride did not produce active reversal of anesthesia as evident by the arousal scores that remained at the levels observed during saline administration under propofol anesthesia. Arousal Score: 1 – EEG activation, 2 – coordinated movements, 3 –attempted return of righting reflex, 4 – return of righting reflex. **C.** Intravenous administration of lisuride did not facilitate passive emergence, i.e., as compared to the saline infusion, there was no significant change in the time to return of righting reflex (RORR) in the post-anesthetic period. For **B-C**, significance symbols (*p* < .05) show post hoc pairwise tests between conditions that were performed with single-step correction for multiple comparisons via Tukey**’s** test: *compared to saline infusion, #compared to DOI infusion. Exact *p* values are reported in the results section and supplementary table S23. **D.** Averaged cohereograms illustrating the global coherence across frequencies (0.1-165 Hz) before, during, and after propofol anesthesia during which the rats received infusion of DOI or lisuride. The black arrow indicates the time of DOI/lisuride delivery. Warmer colors indicate higher coherence while cooler colors indicate less coherence. **E-F.** Line-symbol plots showing global normalized coherence (**E**) and feedback connectivity (measured via normalized symbolic transfer entropy, **F**) for each rat within the theta (4-10 Hz) and high gamma (125-165 Hz) bands during wake, propofol anesthesia, post-drug administration (saline, DOI, or lisuride) during propofol anesthesia, and post-anesthetic recovery after the return of righting reflex. Each data point (pink circle – wake; blue square – anesthesia; orange triangle – anesthesia + saline, DOI, or lisuride; green diamond – post-anesthetic recovery) shows the global normalized coherence (**E**) or feedback connectivity (**F**) for individual rats. For **E-F**, significance symbols (*p* < .05) show post hoc pairwise tests between states that were performed with single-step correction for multiple comparisons via Tukey’s test: *compared to wake, #compared to anesthesia only, §compared to saline or drug (DOI or lisuride) administration during anesthesia. Exact *p* values are reported in the results section and supplementary tables S26-S27 and S30-31. **G.** Topographic maps averaged across all rats illustrate the node degree for each electrode during wake, propofol, post-drug administration (saline, DOI, or lisuride) during anesthesia, and post-anesthetic recovery after the return of righting reflex. Warmer colors indicate higher node degree whereas cooler colors indicate lower node degree. Each white circle represents a single electrode, with 30 electrodes plotted on each topographic map. **H.** Line-symbol plots showing average node degree across electrodes for each rat within the theta and high gamma bands during wake, propofol anesthesia, post-drug administration (saline, DOI, or lisuride) during propofol anesthesia, and post-anesthetic recovery after the return of righting reflex. Each data point represents the average node degree for an individual rat and significance symbols (*p* < .05) show post hoc pairwise tests between states that were performed with single-step correction for multiple comparisons via Tukey**’s** test: *compared to wake, #compared to anesthesia only, §compared to saline or drug (DOI or lisuride) administration during anesthesia. Exact *p* values are reported in the results section and supplementary tables S32-33.

The time to passive emergence was similar between lisuride and saline groups (*p* = .078), but the passive emergence time was longer after lisuride infusion as compared to that after DOI infusion (*p* = .033). Unlike the previous cohort of rats, we did not see a decrease in the time to passive emergence following DOI (compared to saline, *p* = .58) (**Fig. 3C**). This is likely due to our smaller sample size, where 4/9 rats (all male) receiving DOI were unable to be assessed for passive emergence, as they could not be returned to the supine position due to sustained DOI-induced behavioral arousal when it was time to end the anesthetic (see *Experimental Design* in Materials and Methods).

Across conditions, there was no effect of sex on arousal scores (*p* = .053). However, there was a difference in passive emergence time between female and male rats across conditions (*p* = .015), such that female rats took longer on average to regain the righting reflex. This was likely a result of only female rats being assessed for passive emergence in the DOI condition due to continued heightened arousal in the four male rats, which prevented assessment of passive emergence.

### Intravenous Administration of Lisuride during General Anesthesia Did Not Restore High Gamma Non-Directional or Directional Functional Connectivity, or Node Degree

Continuous cohereograms illustrating changes in coherence before and after DOI or lisuride administration during propofol anesthesia are provided in **Fig. 3D**. As with the first cohort, theta global coherence decreased across conditions during propofol anesthesia (*p* < .0001) and as expected, saline infusion did not restore theta coherence (*p* < 1.00). Intravenous infusion of lisuride also did not produce any statistical change in global theta coherence during propofol anesthesia, which remained disrupted and at the level observed during anesthesia only (*p* < 1.00); infusion of DOI decreased theta global coherence even further (*p* = .00063). Theta global coherence remained decreased across conditions compared to baseline wakefulness during the post-propofol recovery period (*p* < .0001) (**Fig. 3E**).

Expectedly, high gamma global coherence decreased across conditions during propofol anesthesia (*p* < .0001), remained low after saline infusion (*p* = 1.00), and recovered to waking levels only during the post-propofol recovery period (*p* = .070). Infusion of DOI during propofol anesthesia increased high gamma coherence to above waking levels (*p* = .013), which returned to baseline waking levels in the post-propofol recovery period (*p* = .38). In contrast, infusion of lisuride decreased high gamma global coherence during propofol anesthesia (*p* = .024), which remained decreased compared to baseline wake state during the post-propofol recovery period (*p* < .0001) (**Fig. 3E**). The effect of lisuride on relative spectral power is illustrated in **Fig. S1E-F.**

High gamma feedback connectivity during propofol anesthesia decreased across all three experimental groups (i.e., infusion of saline, DOI, and lisuride) (*p* < .0001). Infusion of DOI increased high gamma feedback connectivity during propofol anesthesia to that observed during baseline wake state (for DOI vs. anesthesia, *p* < .0001; for DOI vs. wake, *p* = .12). In contrast, neither saline nor lisuride infusion altered high gamma feedback connectivity during propofol anesthesia (*p* ≤ 1.00). As compared to baseline wake state, the post-propofol recovery period was characterized by disrupted high gamma feedback connectivity across all three experimental groups (for saline, *p* < .0001; for DOI, *p* = .00014; for lisuride, *p* = .00029) (**Fig. 3F**). Data for feedforward connectivity are provided in **Fig. S1G** and in the supplementary section.

Topographic maps in **Fig. 3G** depict the average node degree across rats for each electrode within each state, frequency band, and anesthetic for each condition. Theta node degree decreased during propofol anesthesia across the experimental groups (for saline, *p* = .00055; for DOI, *p* = .00012; for lisuride, *p* < .0001) and decreased even further after the infusion of DOI (*p* = .00081) but not saline or lisuride infusion (*p* < 1.00). Theta node degree remained decreased (compared to wake) across experimental groups during the post-propofol recovery period (for saline, *p* = .016; for DOI, *p* = .0051; for lisuride, *p* = .00015 (**Fig. 3H**).

Similar to that observed with theta node degree, high gamma node degree also decreased during propofol anesthesia across the experimental groups (*p* < .0001). Infusion of DOI during propofol anesthesia increased high gamma node degree to waking levels (for DOI vs. anesthesia, *p* < .0001; for DOI vs. wake, *p* = .12), which remained elevated during the post-propofol recovery period (*p* = .43). There was no statistical change in high gamma node degree after the infusion of saline or lisuride (for saline: *p* = 1.00; for lisuride: *p* = .34). High gamma node degree returned to waking levels during the post-propofol recovery period after the saline infusion (*p* = .13) but not after the infusion of lisuride (*p* < .0001) (**Fig. 3H**).

## Discussion

In this study, we report three primary findings. First, we show that 2,5-dimethoxy-4-iodopamphetamine (DOI), a serotonergic psychedelic, can reverse the state of general anesthesia induced by distinct anesthetic agents and that the behavioral arousal was accompanied by increases in neurophysiologic coherence, directed connectivity, and node degree in the high gamma band to the levels observed during baseline wake state. Second, the effect of DOI on anesthetic reversal and restoration of brain dynamics is blocked by a 5-HT2A antagonist. Third, intravenous administration of lisuride, a non-psychedelic 5-HT2A agonist, failed to produce either anesthetic reversal or restoration of brain dynamics as observed after the administration of the psychedelic 5-HT2A agonist, DOI. Previous studies have shown that pharmacological or electrical stimulation of different brain areas can induce waking behavior in anesthetized animals.^1,18,33–40^ Among the systemically delivered drugs, dopaminergic psychostimulants has been shown to induce active emergence whereas caffeine can accelerate passive emergence from anesthesia,^2–6,41^ but neither has been shown to restore the neurophysiological indices associated with normal wakefulness beyond spectral power. In a previous study from our lab, we showed that cholinergic stimulation of medial prefrontal cortex induced wake-like behavior during continued delivery of sevoflurane anesthesia.^18^ However, the active reversal of sevoflurane anesthesia was not accompanied by the restoration of functional connectivity, which remained disrupted despite the presence of wake-like behavior. In contrast to these previous studies with psychostimulants and cholinergic agonism, we demonstrate that the serotonergic psychedelic DOI can reverse general anesthesia and simultaneously restore network features typically observed during wakefulness and cognition.^29,42–44^ Thus, the use of EEG measures beyond spectral changes, particularly at higher frequencies, allows us to evaluate arousal and make inferences about the level of cognition and consciousness.

Our results show restoration of non-directional and directional functional connectivity and functional brain network structure in the gamma—but not theta—band during active emergence from general anesthesia. Previous literature has noted that feedback and feedforward information flow is disrupted in the theta and higher gamma bands during slow-wave sleep and general anesthesia in rats.^29^ However, theta connectivity is restored during rapid eye movement sleep while higher gamma connectivity remains suppressed,^29^ suggesting that activation within higher gamma frequencies may better track changes in level of arousal. This may explain why DOI-induced emergence during general anesthesia was accompanied by a restoration of EEG measures within high gamma, but not theta, band. Notably, EEG measures within the theta band decreased following infusion of DOI during propofol or isoflurane anesthesia. It is worth mentioning that previous studies have reported similar decreases in power and functional connectivity within lower frequencies (1-40 Hz) in rats following the administration of serotonergic psychedelics during wakefulness.^45,46^

We also show that DOI’s effects on behavioral arousal and brain activity were completely blocked by pretreatment with the 5-HT2A antagonist volinanserin, providing evidence for the direct involvement of 5-HT2A receptors, which have been reported as the primary mediators of psychedelic effects associated with serotonergic psychedelics.^8^ Additionally, this is consistent with the arousal-promoting effects of serotonergic psychedelics as reported in previous rodent sleep studies and a study on passive emergence from isoflurane anesthesia.^25–28,47^ Of note, as opposed to the infusion of the psychedelic 5-HT2A agonist DOI, the infusion of lisuride, a non-psychedelic 5-HT2A partial agonist, produced neither behavioral arousal nor EEG activation during propofol anesthesia. Lisuride is frequently used in psychedelic studies as a non-psychedelic serotonergic control given its lack of psychoactive effects in humans and high affinity for the 5-HT2A receptor.^48–52^ Thus, the differential effects of DOI and lisuride—both 5-HT2A agonists—on anesthetic emergence suggest that the downstream effects specific to DOI are likely involved in inducing behavioral arousal.

This study has several limitations. First, we only tested the ability of one serotonergic psychedelic, DOI, to induce arousal during general anesthesia. Thus, it remains to be tested whether other serotonergic psychedelics, such as psilocybin, lysergic acid diethylamide or N, N-dimethyltryptamine can also be used to reverse general anesthesia. We also did not investigate the role of 5-HT2C receptors in this study, which may be of interest given that DOI is a potent agonist at both 5-HT2A and 5-HT2C receptors.^53^ Lastly, despite the restoration of function connectivity and functional brain network organization during behavioral recovery, our data cannot speak to the level of cognition or awareness of the rats during DOI-induced active emergence from general anesthesia. Despite these limitations, our data demonstrate that DOI can reverse the state of general anesthesia induced by two molecularly, pharmacologically, and mechanistically different anesthetics and that the reversal of general anesthesia is accompanied by the restoration of EEG surrogates of information transfer and functional network architecture that are often associated with normal wakefulness. While these findings are mediated by the 5-HT2A receptor, they appear to be associated with psychedelic rather than non-psychedelic agonists. Collectively, these results provide support for translational study of serotonergic psychedelics as a reversal agent for general anesthesia and as a therapeutic agent for pathological states of unconsciousness.^54,55^

## Materials and Methods

All experiments were approved by the Institutional Animal Care and Use Committee at the University of Michigan and were conducted in accordance with the Guide for the Care and Use of Laboratory Animals (National Academies Press, 8th Edition, Washington DC, 2011) and ARRIVE Guidelines. Adult Sprague Dawley rats (n=38, 17 male and 21 female, 300-500 g, Charles River Laboratories) were individually housed with *ad libitum* food and water in a temperature-controlled facility and maintained on a 12 hour:12 hour light:dark cycle (lights ON at 8:00 a.m.).

### Surgical Procedures

Rats were anesthetized with 4-5% isoflurane in 100% oxygen in an air-tight clear rectangular chamber (10.0 inch. x 4.8 inch. x 4.2 inch). Following anesthetic induction, the head and the area over the jugular vein in the neck were shaved, and the rats were immobilized in a stereotaxic frame using blunt ear bars (Model 963, David Kopf Instruments, Tujanga, CA). Isoflurane was delivered via a rat nose cone (Model 906, David Kopf Instruments, Tujunga, CA) for the duration of the surgery and was titrated (1-2%) to maintain the absence of the pedal withdrawal reflex, the presence of regular breathing pattern, and a capillary refill time of < 2 seconds in the extremities, which were monitored every 15 minutes. The delivered anesthetic concentration was monitored continuously using an anesthetic agent analyzer (Capnomac Ultima, Datex Medical Instrumentation, Tewksbury, MA). The rectal temperature was monitored and maintained at 37.0 ± 1 °C using a small animal far-infrared heating pad (RT-0502, Kent Scientific, Torrington, CT). The rats were administered subcutaneous buprenorphine (0.01 mg/kg; Buprenex, Par Pharmaceutical, Chestnut Ridge, NY; NDC 42023-179-05) and carprofen (5 mg/kg; Zoetis; NADA #141-199) for presurgical analgesia and cefazolin (20 mg/kg; West-Ward-Pharmaceutical, Eatontown, NJ; NDC 0143-9924-90) as a prophylactic antibiotic. A mid-sagittal incision was made to expose the cranial surface. Thirty burr holes were drilled across the cranial surface for implantation of custom-made stainless steel screw electrodes to record electroencephalogram (EEG). The EEG electrodes were implanted in a regularly spaced grid of eight rows, parallel to the coronal suture, and between 4 mm anterior and 10 mm posterior to Bregma. Each column of electrodes was parallel and lateral (2 mm to 4.5 mm) to the mid-sagittal suture. Stainless steel screws were also implanted over the nasal sinus and cerebellum to serve as a reference and ground electrode, respectively. Additionally, rats were surgically fitted with an indwelling catheter (MRE-040, Micro-Renathane tubing, Braintree Scientific, Braintree, MA) in the jugular vein for intravenous infusion of DOI (0.5 mg/kg), 0.9% saline, propofol (600-1000 µg/kg/min), volinanserin (25 µg/kg), and/or lisuride (0.05 mg/kg). The free end of electrode wires was secured to a 32-pin Mill-Max connector (Mouser Electronics, Mansfield, TX) and the entire assembly (including the catheter) was affixed to the cranial surface using dental acrylic (Cat No. 51459, Stoelting Co, Woodlake, IL). Post-surgical analgesia was achieved via subcutaneous buprenorphine (0.03 mg/kg) administered every 8-12 hours for 48 hours. The rats were provided at least 7-10 days of post-surgical recovery prior to the start of experiments.

### Pharmacological Agents

Propofol, isoflurane, and 0.9% sterile saline were all procured from the University of Michigan. DOI was obtained from Cayman Chemical (Ann Arbor, MI) and was dissolved in 1 mL of 0.9% sterile saline to create a 5 mg/mL solution. Lisuride was obtained from Tocris Bioscience (Bio-Techne Corporation, Minneapolis, MN) and was mixed with 1 mL of dimethyl sulfoxide (DMSO) to create a 10 mg/mL solution. The same injection volume (500 µL) and flow rate (1-minute infusion using an automated pump) were used for saline, DOI (0.5 mg/kg), and lisuride (0.05 mg/kg) injections. Volinanserin was obtained from Cayman Chemical (Ann Arbor, MI) and was dissolved in 535.475 µL of DMSO to create a 1.87 mg/mL solution, which was delivered as a manual intravenous bolus (25 µg/kg) over 10 seconds.

### Experimental Design

The rats were conditioned to being handled by the experimenters and habituated to the EEG recording setup/chamber for at least one week prior to the start of experiments. In a random order, each rat underwent up to six different experiments with a 5-7 day washout period between the consecutive experiments: i) propofol + DOI, n=10 male and 11 female, ii) propofol + saline, n=10 male and 11 female, iii) propofol + volinanserin + DOI, n=7 male and 9 female, iv) isoflurane + DOI, n=5 male and 9 female, v) isoflurane + saline, n=5 male and 6 female, and vi) isoflurane + volinanserin + DOI, n=5 male and 6 female. On the day of the experiment, rats were connected to the EEG recording system between 9:00 a.m.–11:00 a.m. To minimize handling effects, the baseline EEG data collection started 30-60 minutes after connecting the rats to the EEG recording system and continued for 20 minutes during which we attempted to maintain a constant arousal state by gentle tapping on the cage or introducing novel stimuli if the rats appeared drowsy and/or the EEG showed slow waves indicative of drowsiness. After 20 minutes of baseline data collection, the rats were anesthetized with either propofol or isoflurane. For propofol, an intravenous infusion was started at 1000 µg/kg/min and maintained for 20 minutes, after which the infusion was continued at a lower concentration (600-650 µg/kg/min) for an additional 20 minutes. This dosing regimen was necessitated because continuous intravenous infusion of general anesthesia at a high dose will eventually produce anesthetic overdose and death, if not down-titrated over time. The propofol dose was based on similar previous experiments^29^ conducted in our laboratory and preliminary experiments in which we titrated the dose to achieve a consistent loss of righting reflex and high voltage EEG, indicative of an unconscious state. For isoflurane administration, the rats were briefly disconnected from the EEG system and placed in an anesthetic induction chamber (10.0 inches × 4.8 inches × 4.2 inches) with a continuous inflow of 2.5% isoflurane (2 L/min). Upon the loss of righting reflex, the rats were immediately placed in the experimental chamber and reconnected to the EEG system for the remainder of the experiment where anesthesia was maintained with isoflurane (1.1-1.5%) for 40 minutes. After 40 minutes of propofol or isoflurane anesthesia, the rats received i) 0.5 mg/kg DOI, ii) 0.9% saline, or iii) 0.5 mg/kg DOI preceded by 25 µg/kg volinanserin, which was delivered five minutes prior to the DOI infusion. The effect on behavioral arousal was measured using a behavioral arousal score (see *Arousal Scoring Criteria* for details) in which a score of ‘0’ indicated no change in arousal levels at all and a score of ‘4’ represented the return of righting reflex (RORR), which is a surrogate for recovery from anesthesia in rodents. The changes in behavioral arousal were assessed within the first 5 minutes after the drug/saline infusion. Our primary behavioral outcome measure was active emergence from anesthesia (i.e., arousal in the presence of anesthesia) after DOI administration because of which general anesthesia continued uninterrupted until 20 minutes after drug/saline delivery. Additionally, to determine the time to passive emergence (i.e., time to RORR after the cessation of anesthetic delivery) as a secondary behavioral outcome measure, the rats which showed active emergence were returned to a supine position during the last 5 minutes of anesthetic delivery. After the RORR in the post-anesthetic period, the data collection continued for an additional 20 minutes. To determine the effects of lisuride—a non-psychedelic 5-HT2A partial agonist—on arousal and brain activity during propofol anesthesia, we conducted the following sets of experiments in a separate cohort of rats (n=7 male and 6 female): i) propofol + DOI, n=4 male and female, ii) propofol + saline, n=4 male and 5 female, and iii) propofol + lisuride, n=7 male and 6 female. The experimental procedures, including infusion of lisuride (0.05mg/kg, 500 µL over 1 minute), were identical to those described above for DOI.

The EEG data were continuously acquired during the entire duration of the experiment, except during isoflurane induction. The experimental sessions were video recorded starting 10 minutes before the infusion of drug/saline. Oxygen saturation, heart rate, and respiration rate were monitored using a pulse oximetry sensor (MouseOx, Starr Life Science Corp., Oakmont, PA) positioned around the neck. The core body temperature was maintained at 37 ± 1 ◦C using a rectal probe (RET-2 ISO, Physitemp Instruments, Inc., Clifton, NJ) connected via a feedback temperature controller (TCAT-2LV, Physitemp Instruments, Inc., Clifton, New Jersey) to a small animal heating pad (Kent Scientific Co., Torrington, Connecticut).

### Arousal Scoring Criteria

To quantify changes in level of arousal after the drug/saline infusion during general anesthesia (i.e., active emergence), an experimenter blinded to the drug condition reviewed and scored each video to determine rat behavior during the first 5 minutes after the end of the drug/saline infusion. The scoring criteria were as follows: 1=EEG activation, 2=coordinated/purposeful movements (e.g., lifting head, pawing, sniffing.), 3=attempts at RORR, and 4=RORR. The blinded reviewer also assessed each video for the time to RORR following the end of the anesthetic delivery (i.e., passive emergence; see *Experimental Design* for more details).

### EEG Data Acquisition and Analyses

Monopolar EEG data were acquired using a 32-channel head stage and one of two different recording systems: a Cereplex Direct system paired with the Cereplex Direct Software Suite (signals digitized at 1000 Hz; Blackrock Microsystems, Salt Lake City, UT) or a PZ5 NeuroDigitizer amplifier paired with Synapse software (signals digitized at 1017 Hz; Tucker-Davis Technologies Inc, Alachua, FL). All data analyses were performed using MATLAB (version 2022b; MathWorks, Inc.; Natick, MA). The raw EEG waveform and spectrogram for each file were visually inspected in 9-second windows for movement artifacts or noisy data due to hardware issues. The EEG data segments during the baseline or post-anesthesia recovery wakefulness periods that contained slow waves indicative of sleep or a drowsy state were excluded from analyses. Data were recleaned in 3-second windows following the drug infusion period to remove segments containing the head twitch response. Bad channels were also identified and excluded from analysis. Subsequent analyses utilized clean data segments (total lengths 54-297 seconds, or roughly 1-5 minutes) from the following states: i) Wake, i.e., up to 20 minutes before the start of anesthesia, ii) Anesthesia, i.e., 5 minutes before DOI, lisuride, or saline infusion during propofol or isoflurane anesthesia, iii) Drug/Saline infusion during anesthesia, i.e., 5 minutes after the end of the DOI, lisuride, or saline infusion during propofol or isoflurane anesthesia, and iv) Post-anesthesia Recovery Wake, i.e., the first 15 minutes after the anesthetic had been turned off and the RORR. Rats with less than 54 seconds of data for a given state were excluded from analyses for only that state. Measures of power spectral density, magnitude-squared coherence, normalized symbolic transfer entropy, and node degree were computed for each state within the theta (4–10 Hz) and high gamma (125–165 Hz) bands. We focused on these frequency bands because previous studies have shown a clear relationship between these frequency ranges and states of arousal in rodents.^29,30^

### Power Spectral Density (PSD) Analysis

Welch’s method (pwelch.m) was used to calculate PSD at the channel level for 0.5-200 Hz for each state (9-second moving windows; 3-second hamming window). Relative spectral power was calculated as the absolute spectral power within a given frequency divided by the total absolute spectral power across all frequencies and then averaged across all time windows for each state. Finally, relative power values were averaged across all frequencies within each band and across all channels for the global relative power within each frequency band.

### Magnitude-squared Coherence Analysis

Magnitude-squared coherence (mscohere.m) was computed for 0.5-200 Hz across all channel pairs using 9-second non-overlapping windows. Coherence values for each channel pair were normalized using the mean and standard deviation of surrogate datasets (n=50) generated through phase shuffling while preserving the amplitude distribution of each signal. The resulting Z-scores were averaged across all windows and channel pairs for each frequency band in each state, excluding pairs with bad channels.

### Normalized Symbolic Transfer Entropy Analysis

Normalized symbolic transfer entropy (NSTE) is a measure of directional functional connectivity and is a surrogate measure of information exchange rooted in information theory that allows one to evaluate interactions between different electrodes. It is non-parametric, nonlinear, and model-free. At its core, NSTE examines the inferred causal influence of source signal X on target signal Y by comparing two information values to answer the following question: is the ability to predict the future of target signal Y improved by knowing the past of signals X and Y? If so, X is inferred to have a causal influence on Y. The precise methodology for computing NSTE is outlined in previous human^56–58^ and rat studies from our group.^29,30,59–61^ To reduce the computational burden, data were resampled from 1000 Hz/1017 Hz to 500 Hz/678 Hz prior to calculating NSTE. Data were then filtered using a fourth-order Butterworth filter (butter.m and filtfilt.m, MATLAB Signal Processing Toolbox). We computed NSTE values for ipsilateral pairs of frontal (electrode no. 6 and 7) and parietal channels (electrode no. 18 and 19) to assess feedforward (parietal-to-frontal) and feedback (frontal-to-parietal) connectivity for each frequency band within each state (i.e., Wake, Anesthesia, Drug/Saline infusion during anesthesia, and Post-anesthesia Recovery Wake). Our selection of electrode pairs was based on the pairs of channels demonstrating the highest NSTE values during wakefulness, which was consistent with our prior publications.^29,30,59–61^ Following averaging across time windows for each state, NSTE values within each frequency band for both pairs of electrodes were averaged together for statistical comparison.

### Node Degree Analysis

Node degree was calculated using binarized data achieved through thresholding of the normalized coherence values. After averaging Z-scores across time for each state, the resulting matrix was binarized using a threshold equal to the Z-score for alpha = 0.01 (i.e., Z = 2.3263), where Z-scores above the threshold were replaced with a “1” and Z-scores below the threshold were replaced with a “0”. The resulting binarized matrix was used to calculate the node degree (i.e., number of connections or “1s”) within each frequency band for each channel using the Brain Connectivity Toolbox (degrees_und.m).^62^ For each frequency band, node degree values were averaged across channels for statistical analyses.

### Statistical Analyses

Statistical analyses were completed using R software^63^ (RStudio 2020 version 4.0.0) and the packages emmeans and lme4.^64,65^ Sample sizes were based on a priori power analyses using behavioral arousal data from pilot studies in our laboratory and previous publications.^18,30–32^ Our primary outcome measures included i) arousal score, i.e., active emergence, ii) the time to RORR following discontinuation of propofol or isoflurane anesthesia, i.e., passive emergence, and iii) the average relative spectral power, global normalized coherence, feedback and feedforward NSTE, and average node degree for each frequency band. A linear mixed model was used to compare the arousal score and time to passive emergence across experimental groups (saline vs. DOI vs. DOI + volinanserin; saline vs. DOI vs. lisuride), where “condition” and “sex” were treated as fixed factors and “subject” as a random intercept. An alpha threshold of *p* < 0.05 was used and Tukey’s method was applied to adjust *p* values for the number of contrasts between conditions. For each EEG measure, a linear mixed model was also implemented, where “condition,” “state,” and “sex” were treated as fixed factors and “subject” as a random intercept. We then conducted a likelihood ratio test to determine whether our model was improved by including condition as an interaction term. Results of pairwise comparison tests between each state within each condition are reported in the text for measures where including “condition” as an interaction term improved the model (*p* < 0.05 for likelihood ratio test), using an alpha threshold of *p* < 0.05 and Tukey’s method to correct for the number of contrasts across states (Wake, Anesthesia, Drug/Saline infusion during anesthesia, Post-anesthesia Recovery Wake). P values for each pairwise comparison are reported in the text and the corresponding degrees of freedom, t-statistic, and 95% confidence interval (CI), as well as the results from each likelihood ratio test, are reported in **tables S1-S33**. The results from likelihood ratio tests and sex effects are reported in the supplementary section.

