## Supplementary Materials for "Psychedelic-mediated Reversal of General Anesthesia and Restoration of Brain Dynamics in Rat"

**The PDF file includes:**

Supplementary Text
Fig. S1 and legend
Tables S1 to S33 and legends
Legends for Movies S1 and S2

**Other Supplementary Materials for this manuscript include the following:**

Movies S1 and S2

### Supplementary Text

#### *Intravenous Administration of DOI Restored High Gamma, but not Theta, Relative Power*

Representative spectrograms showing relative power before and after DOI or DOI following pretreatment with volinanserin during propofol or isoflurane anesthesia are illustrated in **Fig. S1A-B**. Likelihood-ratio tests revealed that including condition (i.e., saline, DOI, or DOI following pretreatment with volinanserin) as an interaction term led to a better model fit for relative power in the theta (for propofol, $p = .00087$; for isoflurane, $p = .0097$) and high gamma (for propofol, $p < .0001$; for isoflurane, $p = .00019$) band. There was no difference in theta (for propofol, $p < 1.00$; for isoflurane, $p = .10$) or high gamma (for propofol, $p < 1.00$; for isoflurane, $p = .34$) relative power between female and male rats across conditions. Subsequent pairwise comparison tests revealed that, compared to wake, theta relative power was significantly decreased across conditions during propofol or isoflurane anesthesia ($p < .0001$). DOI infusion decreased theta relative power even further during either anesthetic (for propofol, $p = .036$; for isoflurane, $p = .0087$). There was no change following the saline infusion ($p = 1.00$) and the effect of DOI on theta relative power was blocked by pretreatment with volinanserin (p < 1.00). Theta relative power remained decreased (compared to wake) across conditions following the cessation of propofol ($p < .0001$) or isoflurane anesthesia (for saline, $p = .00018$; for DOI, $p < .0001$; for DOI + volinanserin, $p = .00021$) (**Fig. S1C**).

High gamma relative power decreased following propofol or isoflurane anesthesia across conditions ($p < .0001$); the power increased following the infusion of DOI during both anesthetics ($p < .0001$) but remained decreased compared to waking levels (for propofol, $p < .0001$; for isoflurane, $p = .0089$). This effect was blocked by pretreatment with volinanserin for both anesthetics ($p = 1.00$) and did not occur following saline infusion ($p = 1.00$). In the DOI condition, high gamma relative power increased even further following the cessation of either anesthetic (for propofol, recovery vs. DOI: $p < .0001$; for isoflurane, recovery vs. DOI: $p = .00035$) and was elevated compared to wake in propofol experiments ($p = .00073$). High gamma relative power increased during the recovery period following the end of propofol or isoflurane anesthesia for both control conditions ($p < .0001$) (**Fig. S1C**).

***Extended Results: Intravenous Administration of DOI Restored High Gamma, but not Theta, Non-Directional Connectivity***

Likelihood-ratio tests revealed an improved model fit by including condition (i.e., saline, DOI, or DOI following volinanserin pretreatment) as an interaction term for both frequency bands and anesthetics ($p < .0001$). There was no difference in global normalized coherence values between female and male rats across conditions for the theta (for propofol, $p < 1.00$; for isoflurane, $p = .30$) or high gamma (for propofol, $p = .38$; for isoflurane, $p < 1.00$) band.

***Extended Results: Intravenous Administration of DOI Restored High Gamma, but not Theta, Directional Connectivity***

For both feedback and feedforward connectivity, the results of each likelihood-ratio test indicated that including condition (i.e., saline, DOI, DOI following volinanserin pretreatment) as an interaction term improved model fit for the high gamma band for propofol or isoflurane experiments ($p < .0001$), as well as for the theta band for propofol experiments (for feedforward, $p < .0001$; for feedback, $p = .0037$). Theta feedforward ($p = .018$) and feedback ($p = .029$) connectivity were lower in female rats compared to male rats across conditions in isoflurane experiments but similar between female and male rats for propofol experiments (for feedforward, *p* < 1.00; for feedback, *p* = .36). High gamma feedforward (for propofol, *p* < 1.00; for isoflurane, *p* = .28) and feedback (for propofol, *p* = .24; for isoflurane, *p* = .10) connectivity were similar between female and male rats across conditions.

Theta feedforward connectivity decreased during propofol anesthesia across conditions (p < .0001). DOI infusion further decreased feedforward connectivity during propofol anesthesia (p < .0001); this decrease was blocked by pretreatment with volinanserin (p = 1.00) and did not occur following the saline infusion (p = .84). Theta feedforward connectivity remained decreased during the post-propofol recovery period across conditions (saline: p < .0001; DOI: p = .00086; DOI + volinanserin: p < .0001) (**Fig. S1D**).

High gamma feedforward connectivity decreased across conditions during propofol or isoflurane anesthesia (*p* < .0001). DOI infusion increased high gamma feedforward connectivity to above waking levels during either anesthetic (for propofol, *p* = .0026; for isoflurane, *p* = .00058). Following DOI infusion, feedforward high gamma connectivity remained at or above baseline wake during the post-anesthesia recovery period (for propofol, *p* = .00051; for isoflurane, *p* = .29). There was no statistical change in high gamma feedforward connectivity during either anesthetic following saline infusion (*p* < 1.00) or DOI infusion after pretreatment with volinanserin (*p* = 1.00) and, compared to wake, high gamma feedforward connectivity for both control conditions remained decreased during the post-propofol (for saline, *p* < .0001; for DOI + volinanserin, *p* = .022) and post-isoflurane (for saline, *p* = .067; for DOI + volinanserin, *p* = .012) recovery period (**Fig. S1D**).

***Extended Results: Intravenous Administration of DOI Restored High Gamma, but not Theta, Node Degree***

Likelihood-ratio tests confirmed that the inclusion of condition (i.e., saline, DOI, DOI + volinanserin) as an interaction term improved the model fit for both frequency bands and anesthetics (all $p < .0001$). Node degree values were similar between female and male rats across conditions for the theta (for propofol and isoflurane, $p < 1.00$) and high gamma (for propofol, p = .17; for isoflurane, $p < 1.00$) band.

#### ***Intravenous Administration of Lisuride during General Anesthesia Did Not Restore High Gamma Relative Power***

Representative spectrograms showing relative spectral power before and after DOI or lisuride during propofol anesthesia are illustrated in **Fig. S1E.** Likelihood-ratio tests revealed that including condition (i.e., saline, DOI, lisuride) as an interaction term led to a better model fit for relative power in the theta ($p = .00056$) and high gamma ($p = .00033$) band during propofol anesthesia. Relative power did not different between female and male rats for the theta ($p = .27$) or high gamma ($p = .55$) band. Theta relative power decreased during propofol anesthesia across conditions (for saline, $p = .010$; for DOI, $p = .0015$; for lisuride, $p = .00014$). Lisuride infusion during propofol anesthesia increased theta relative power ($p = .011$) while there was no statistical change after saline infusion ($p < 1.00$). In this new cohort of rats, infusion of DOI during propofol anesthesia did not decrease theta relative power ($p = .43$). Across conditions, theta relative power remained decreased during the post-propofol recovery period compared to baseline wakefulness (for saline, $p < .0001$; for DOI, $p = .00058$; for lisuride, $p < .0001$) (**Fig. S1F**).

High gamma relative power decreased across conditions during propofol anesthesia ($p < .0001$). Infusion of DOI, but not lisuride or saline, increased high gamma relative power during propofol anesthesia (for saline, $p = 1.00$; for DOI, $p = .034$; for lisuride, $p = 1.00$). During the post-propofol recovery period, high gamma relative power returned to waking levels in the DOI ($p = .063$) and saline ($p = .072$) groups but not in the lisuride group ($p = .0017$).

***Extended Results: Intravenous Administration of Lisuride during General Anesthesia Did Not Restore High Gamma Non-Directional or Directional Functional Connectivity, or Node Degree***

Model fit was improved by including condition (i.e., saline, DOI, lisuride) as an interaction term for global normalized coherence in the theta ($p = .011$) and high gamma ($p < .0001$) band. There was no difference in global coherence values between female and male rats across experimental conditions for either of the frequency bands ($p < 1.00$).

Model fit was improved by the inclusion of condition (i.e., saline, DOI, lisuride) as an interaction term for feedback and feedforward connectivity (measured via normalized symbolic transfer entropy) for the high gamma band ($p < .0001$), as well as for feedforward connectivity in the theta band ($p = .0023$). Feedforward and feedback connectivity were similar between female and male rats across conditions for both frequency bands ($p < 1.00$). Theta feedforward connectivity decreased across experimental conditions during propofol anesthesia (for saline, $p = .00023$; for DOI, $p < .0001$; for lisuride, $p < .0001$). Infusion of DOI decreased theta feedforward connectivity even further during propofol anesthesia ($p = .034$), which recovered to waking levels during the post-propofol recovery period ($p = .31$). There was no statistical change following the infusion of saline ($p = 1.00$) or lisuride ($p < 1.00$) during propofol anesthesia and, compared to baseline wake state, theta feedforward connectivity remained decreased in the post-propofol recovery period (for saline, $p = .0039$; for lisuride, $p < .0001$) **(Fig. S1G)**.

High gamma feedforward connectivity decreased across conditions during propofol anesthesia ($p < .0001$). Infusion of DOI increased high gamma feedforward connectivity during propofol anesthesia to waking levels (for DOI vs. anesthesia, $p < .0001$; for DOI vs. wake, $p = .37$). In contrast, neither saline nor lisuride infusion altered high gamma feedforward connectivity during propofol anesthesia (for saline, $p = 1.00$; for lisuride, $p = .53$). High gamma feedforward connectivity across experimental conditions remained decreased during the post-propofol recovery period compared to wake (for saline, $p < .0001$; for DOI, $p = .0061$; for lisuride, $p < .0001$) (**Fig. S1G**).

Likelihood ratio tests revealed an improved model fit for node degree in the theta ($p = .040$) and high gamma band ($p < .0001$) when including condition (i.e., saline, DOI, lisuride) as an interaction term and, across experimental conditions, there was no difference in node degree between sexes for either of the frequency bands ($p < 1.00$).

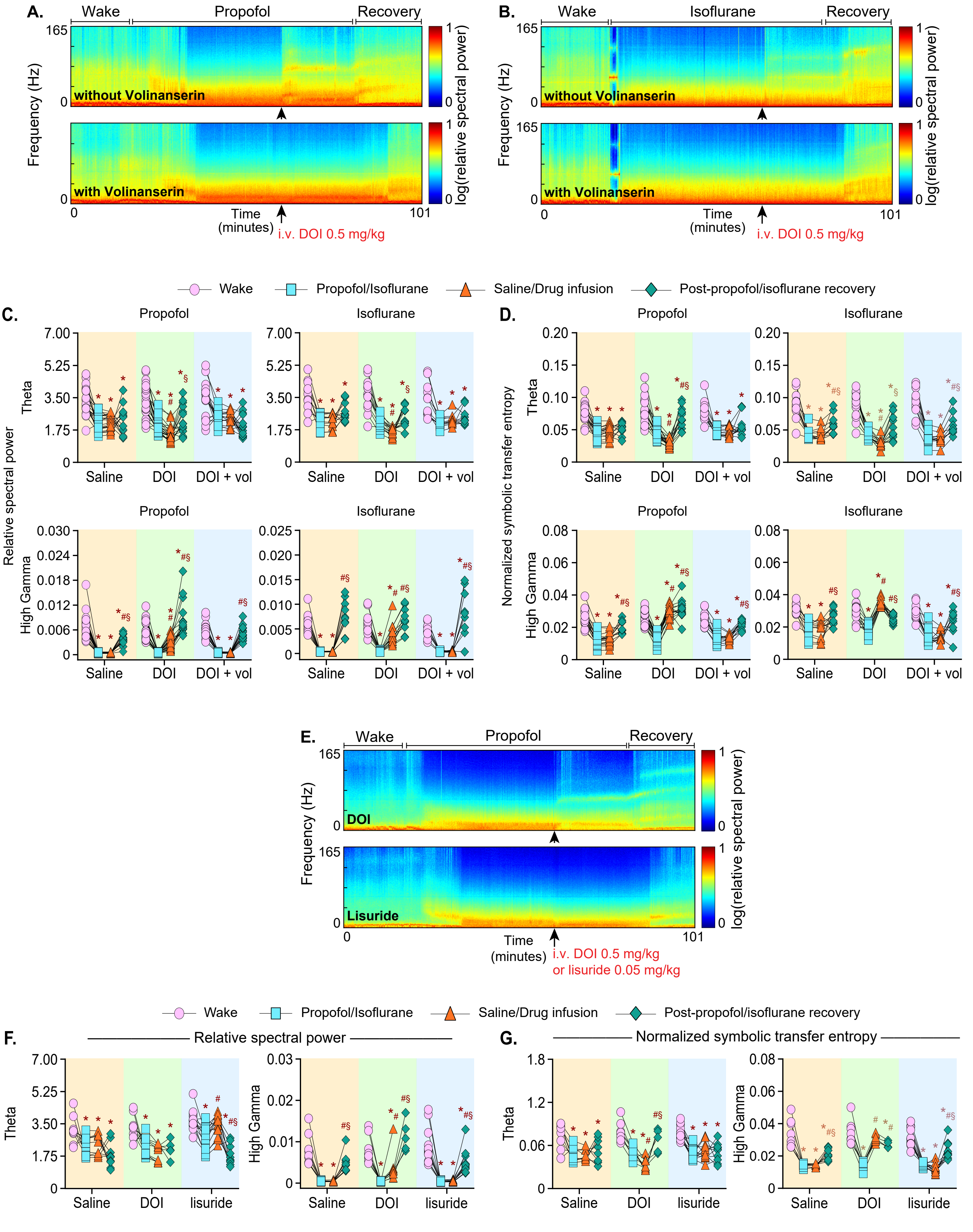
A.
Wake
Propofol
Recovery
165
0
without Volinanserin
with Volinanserin
Frequency (Hz)
log(relative spectral power)
Time
(minutes)
101
i.v. DOI 0.5 mg/kg
B.
Isoflurane
Wake
Post-propofol/isoflurane recovery
Propofol/Isoflurane
Saline/Drug infusion
C.
Relative spectral power
Theta
High Gamma
Saline
DOI
DOI + vol
D.
Normalized symbolic transfer entropy
E.
DOI
Lisuride
i.v. DOI 0.5 mg/kg
or lisuride 0.05 mg/kg
F.
G.
lisuride

**Fig. S1. Relative Spectral Power and Feedforward Connectivity Before and After Saline, DOI, DOI Pretreated with Volinanserin, or Lisuride during Propofol or Isoflurane Anesthesia.**

**A-B**. Spectrograms illustrating the global relative spectral power across frequencies (0.1-165 Hz) before, during, and after propofol (**A**) or isoflurane (**B**) during which the rats received DOI or DOI with volinanserin pretreatment. The black arrow indicates the time of DOI delivery. Warmer colors indicate higher power while cooler colors indicate less power. **C-D.** Line-symbol plots show global relative spectral power (**C**) or feedforward connectivity (measured via normalized symbolic transfer entropy, **D**) within the theta (4-10 Hz) and high gamma (125-165 Hz) bands during wake, anesthesia (propofol or isoflurane), post-drug administration (saline, DOI, or DOI with volinanserin pretreatment during propofol or isoflurane anesthesia), and post-anesthetic recovery after the return of righting reflex. Each data point (pink circle – wake**;** blue square – anesthesia**;** orange triangle – anesthesia + saline, DOI, or DOI with volinanserin pretreatment; green diamond – post-anesthetic recovery) shows the global relative spectral power (**C**) or feedforward connectivity (**D**) for an individual rat. **S**ignificance symbols ($p < .05$) show post hoc pairwise tests between states that were performed with single-step correction for multiple comparisons via Tukey**'s** test**:** *compared to wake, #compared to anesthesia only, §compared to saline or drug (DOI or DOI pretreated with volinanserin) administration during anesthesia. Exact *p* values are reported in the supplementary section and supplementary tables S3-6 and S11-14. **E**. Spectrograms illustrating the global relative spectral power across frequencies (0.1-165 Hz) before, during, and after propofol anesthesia during which rats received DOI or lisuride. The black arrow indicates the time of DOI/lisuride delivery. Warmer colors indicate higher power while cooler colors indicate less power. **F-G.** Line-symbol plots show global relative spectral power (**F**) or feedforward connectivity (measured via normalized symbolic transfer entropy, **G**) within the theta and high gamma bands during wake, propofol anesthesia, post-drug administration (saline, DOI, or lisuride during propofol anesthesia), and post-anesthetic recovery after the return of righting reflex. Each data point (pink circle – wake; blue square – anesthesia; orange triangle – anesthesia + saline, DOI, or lisuride; green diamond – post-anesthetic recovery) shows the global relative spectral power (**F**) or feedforward connectivity (**G**) for an individual rat. **S**ignificance symbols ($p < .05$) show post hoc pairwise tests between states that were performed with single-step correction for multiple comparisons via Tukey**'s** test**:** *compared to wake, #compared to anesthesia only, §compared to saline or drug (DOI or lisuride) administration during anesthesia. Exact *p* values are reported in the supplementary section and supplementary tables S24-25 and S28-29.

**Table S1. Statistical Outputs for Active and Passive Emergence after the Infusion of Saline, DOI, or DOI with Volinanserin Pretreatment during Propofol Anesthesia**

| ***Arousal Score*** | **Estimate** | **df** | ***t*** | ***p*** | **Lower CI** | **Upper CI** |
|---|---|---|---|---|---|---|
| *LMM: Female v. Male* | -0.014 | 18.66 | -0.15 | 0.88 | -0.20 | 0.17 |
| *DOI ~ Saline* | 3.38 | 35.41 | 30.84 | < 0.0001 | 3.11 | 3.65 |
| *DOI + Vol ~ Saline* | 0.00065 | 38.77 | 0.0054 | 1.00 | -0.29 | 0.29 |
| *DOI + Vol ~ DOI* | -3.38 | 38.77 | -28.46 | < 0.0001 | -3.67 | -3.09 |
| ***RORR*** | **Estimate** | **df** | ***t*** | ***p*** | **Lower CI** | **Upper CI** |
| *LMM: Female v. Male* | 242.57 | 16.32 | 2.73 | 0.015 | 69.51 | 414.84 |
| *DOI ~ Saline* | -264.91 | 28.35 | -3.22 | 0.0088 | -468.63 | -61.19 |
| *DOI + Vol ~ Saline* | 100.84 | 27.59 | 1.25 | 0.43 | -98.84 | 300.52 |
| *DOI + Vol ~ DOI* | 365.75 | 28.36 | 4.60 | 0.00023 | 169.12 | 562.39 |

Statistical outputs include the sex effect for the linear mixed model (LMM) and results for each pairwise comparison between conditions (saline, DOI, and DOI with volinanserin pretreatment [DOI + Vol]) for arousal score (i.e., active emergence) and time to the return of righting reflex (RORR) after cessation of the anesthetic (i.e., passive emergence). Corrected p values using Tukey's method are reported for pairwise comparisons. CI = 95% confidence interval; df = degrees of freedom; Estimate = difference in means.

**Table S2. Statistical Outputs for Active and Passive Emergence after the Infusion of Saline, DOI, or DOI with Volinanserin Pretreatment during Isoflurane Anesthesia**

| ***Arousal Score*** | **Estimate** | **df** | ***t*** | ***p*** | **Lower CI** | **Upper CI** |
|---|---|---|---|---|---|---|
| *LMM: Female v. Male* | 0.12 | 10.51 | 0.51 | 0.62 | -0.31 | 0.54 |
| *DOI ~ Saline* | 2.99 | 23.12 | 11.30 | < 0.0001 | 2.32 | 3.65 |
| *DOI + Vol ~ Saline* | 3.31E-16 | 21.15 | 1.20E-15 | 1.00 | -0.70 | 0.70 |
| *DOI + Vol ~ DOI* | -2.99 | 23.12 | -11.30 | < 0.0001 | -3.65 | -2.32 |
| ***RORR*** | **Estimate** | **df** | ***t*** | ***p*** | **Lower CI** | **Upper CI** |
| *LMM: Female v. Male* | -86.09 | 9.00 | -2.07 | 0.069 | -162.39 | -9.78 |
| *DOI ~ Saline* | -126.64 | 20.00 | -2.61 | 0.042 | -249.24 | -4.03 |
| *DOI + Vol ~ Saline* | 166.36 | 20.00 | 3.43 | 0.0071 | 43.76. | 288.97 |
| *DOI + Vol ~ DOI* | 293.00 | 20.00 | 6.05 | < 0.0001 | 170.40 | 415.60 |

Statistical outputs include the sex effect for the linear mixed model (LMM) and results for each pairwise comparison between conditions (saline, DOI, and DOI with volinanserin pretreatment [DOI + Vol]) for arousal score (i.e., active emergence) and time to the return of righting reflex (RORR) after cessation of the anesthetic (i.e., passive emergence). Corrected p values using Tukey's method are reported for pairwise comparisons. CI = 95% confidence interval; df = degrees of freedom; Estimate = difference in means.

**Table S3. Statistical Outputs for Theta Relative Power before and after Infusion of Saline, DOI, or DOI with Volinanserin Pretreatment during Propofol Anesthesia**

| | **Estimate** | **df** | ***t*** | ***p*** | **Lower CI** | **Upper CI** |
|---|---|---|---|---|---|---|
| ***LMM: Female v. Male*** | -0.035 | 18.83 | -0.22 | 0.83 | -0.34 | 0.28 |
| | | **df** | **$X^2$** | ***p*** | | |
| ***Likelihood Ratio Test*** | - | 6.00 | 22.80 | 0.00087 | - | - |
| | **Estimate** | **df** | ***t*** | ***p*** | **Lower CI** | **Upper CI** |
| ***Saline*** | | | | | | |
| *A ~ W* | -1.17 | 187.06 | -6.82 | < 0.0001 | -1.62 | -0.73 |
| *D ~ W* | -1.14 | 187.06 | -6.64 | < 0.0001 | -1.59 | -0.70 |
| *D ~ A* | 0.030 | 187.06 | 0.17 | 1.00 | -0.42 | 0.48 |
| *R ~ W* | -1.17 | 189.13 | -6.02 | < 0.0001 | -1.67 | -0.67 |
| *R ~ A* | 0.0036 | 189.13 | 0.019 | 1.00 | -0.50 | 0.51 |
| *R ~ D* | -0.026 | 189.13 | -0.13 | 1.00 | -0.53 | 0.48 |
| ***DOI*** | | | | | | |
| *A ~ W* | -1.42 | 187.06 | -8.24 | < 0.0001 | -1.86 | -0.97 |
| *D ~ W* | -1.88 | 187.06 | -10.96 | < 0.0001 | -2.33 | -1.44 |
| *D ~ A* | -0.47 | 187.06 | -2.72 | 0.036 | -0.91 | -0.021 |
| *R ~ W* | -1.21 | 188.75 | -6.38 | < 0.0001 | -1.70 | -0.72 |
| *R ~ A* | 0.21 | 188.75 | 1.09 | 0.70 | -0.29 | 0.70 |
| *R ~ D* | 0.67 | 188.75 | 3.55 | 0.0027 | 0.18 | 1.16 |
| ***DOI + Vol*** | | | | | | |
| *A ~ W* | -0.98 | 187.06 | -4.98 | < 0.0001 | -1.49 | -0.47 |
| *D ~ W* | -1.04 | 187.06 | -5.28 | < 0.0001 | -1.55 | -0.53 |
| *D ~ A* | -0.059 | 187.06 | -0.30 | 0.99 | -0.57 | 0.45 |
| *R ~ W* | -1.46 | 187.06 | -7.40 | < 0.0001 | -1.97 | -0.95 |
| *R ~ A* | -0.48 | 187.06 | -2.41 | 0.078 | -0.99 | 0.035 |
| *R ~ D* | -0.42 | 187.06 | -2.11 | 0.15 | -0.93 | 0.094 |

Statistical outputs include the sex effect for the linear mixed model (LMM), the results of the likelihood ratio test, and the results for each pairwise comparison between states of wake (W), anesthesia (A), drug (D), and recovery (R) for saline, DOI, or DOI + volinanserin (DOI + Vol). Corrected p values using Tukey’s method are reported for pairwise comparisons. CI = 95% confidence interval; df = degrees of freedom; Estimate = difference in means.

**Table S4. Statistical Outputs for High Gamma Relative Power before and after Infusion of Saline, DOI, or DOI with Volinanserin Pretreatment during Propofol Anesthesia**

| | **Estimate** | **df** | ***t*** | ***p*** | **Lower CI** | **Upper CI** |
|---|---|---|---|---|---|---|
| ***LMM: Female v. Male*** | -0.00023 | 17.79 | -0.63 | 0.54 | -0.00095 | 0.00048 |
| | | **df** | $X^2$ | ***p*** | | |
| ***Likelihood Ratio Test*** | - | 6.00 | 51.60 | < 0.0001 | - | - |
| | **Estimate** | **df** | ***t*** | ***p*** | **Lower CI** | **Upper CI** |
| ***Saline*** | | | | | | |
| *A ~ W* | -0.0064 | 186.19 | -11.61 | < 0.0001 | -0.0078 | -0.0050 |
| *D ~ W* | -0.0064 | 186.19 | -11.59 | < 0.0001 | -0.0078 | -0.0049 |
| *D ~ A* | 0.000013 | 186.19 | 0.023 | 1.00 | -0.0014 | 0.0014 |
| *R ~ W* | -0.0031 | 189.81 | -5.08 | < 0.0001 | -0.0048 | -0.0015 |
| *R ~ A* | 0.0032 | 189.81 | 5.23 | < 0.0001 | 0.0016 | 0.0048 |
| *R ~ D* | 0.0032 | 189.81 | 5.21 | < 0.0001 | 0.0016 | 0.0048 |
| ***DOI*** | | | | | | |
| *A ~ W* | -0.0058 | 186.54 | -10.39 | < 0.0001 | -0.0072 | -0.0043 |
| *D ~ W* | -0.0032 | 186.54 | -5.82 | < 0.0001 | -0.0047 | -0.0018 |
| *D ~ A* | 0.0025 | 186.19 | 4.63 | < 0.0001 | 0.0011 | 0.0040 |
| *R ~ W* | 0.0024 | 189.69 | 3.91 | 0.00073 | 0.00081 | 0.0040 |
| *R ~ A* | 0.0082 | 189.19 | 13.51 | < 0.0001 | 0.0066 | 0.0098 |
| *R ~ D* | 0.0056 | 189.19 | 9.31 | < 0.0001 | 0.0041 | 0.0072 |
| ***DOI + Vol*** | | | | | | |
| *A ~ W* | -0.0057 | 186.188 | -9.05 | < 0.0001 | -0.0073 | -0.0041 |
| *D ~ W* | -0.0057 | 186.188 | -9.06 | < 0.0001 | -0.0073 | -0.0041 |
| *D ~ A* | -3.5E-06 | 186.188 | -0.0055 | 1.00 | -0.0016 | 0.0016 |
| *R ~ W* | -0.0010 | 186.188 | -1.63 | 0.36 | -0.0027 | 0.00060 |
| *R ~ A* | 0.0047 | 186.188 | 7.42 | < 0.0001 | 0.0030 | 0.0063 |
| *R ~ D* | 0.0047 | 186.188 | 7.42 | < 0.0001 | 0.0030 | 0.0063 |

Statistical outputs include the sex effect for the linear mixed model (LMM), the results of the likelihood ratio test, and the results for each pairwise comparison between states of wake (W), anesthesia (A), drug (D), and recovery (R) for saline, DOI, or DOI + volinanserin (DOI + Vol). Corrected p values using Tukey’s method are reported for pairwise comparisons. CI = 95% confidence interval; df = degrees of freedom; Estimate = difference in means.

**Table S5. Statistical Outputs for Theta Relative Power before and after Infusion of Saline, DOI, or DOI with Volinanserin Pretreatment during Isoflurane Anesthesia**

| | Estimate | df | *t* | *p* | Lower CI | Upper CI |
|---|---|---|---|---|---|---|
| ***LMM: Female v. Male*** | -0.36 | 9.82 | -1.80 | 0.10 | -0.77 | 0.022 |
| | | **df** | $X^2$ | ***p*** | | |
| ***Likelihood Ratio Test*** | - | 6.00 | 16.88 | 0.0097 | - | - |
| | **Estimate** | **df** | ***t*** | ***p*** | **Lower CI** | **Upper CI** |
| ***Saline*** | | | | | | |
| *A ~ W* | -1.38 | 113.20 | -7.09 | < 0.0001 | -1.88 | -0.87 |
| *D ~ W* | -1.38 | 113.20 | -7.10 | < 0.0001 | -1.89 | -0.87 |
| *D ~ A* | -0.0025 | 113.20 | -0.013 | 1.00 | -0.51 | 0.50 |
| *R ~ W* | -0.87 | 113.37 | -4.34 | 0.00018 | -1.39 | -0.35 |
| *R ~ A* | 0.51 | 113.37 | 2.56 | 0.056 | -0.0092 | 1.03 |
| *R ~ D* | 0.51 | 113.37 | 2.57 | 0.054 | -0.0067 | 1.03 |
| ***DOI*** | | | | | | |
| *A ~ W* | -1.49 | 113.20 | -8.65 | < 0.0001 | -1.94 | -1.04 |
| *D ~ W* | -2.05 | 113.20 | -11.88 | < 0.0001 | -2.49 | -1.60 |
| *D ~ A* | -0.56 | 113.20 | -3.23 | 0.0087 | -1.00 | -0.11 |
| *R ~ W* | -1.31 | 115.53 | -7.04 | < 0.0001 | -1.80 | -0.83 |
| *R ~ A* | 0.18 | 115.53 | 0.96 | 0.77 | -0.31 | 0.66 |
| *R ~ D* | 0.73 | 115.53 | 3.94 | 0.00079 | 0.25 | 1.22 |
| ***DOI + Vol*** | | | | | | |
| *A ~ W* | -1.24 | 113.20 | -6.37 | < 0.0001 | -1.74 | -0.73 |
| *D ~ W* | -1.10 | 113.20 | -5.65 | < 0.0001 | -1.60 | -0.59 |
| *D ~ A* | 0.14 | 113.20 | 0.73 | 0.89 | -0.37 | 0.65 |
| *R ~ W* | -0.88 | 113.49 | -4.30 | 0.00021 | -1.42 | -0.35 |
| *R ~ A* | 0.35 | 113.49 | 1.72 | 0.32 | -0.18 | 0.89 |
| *R ~ D* | 0.21 | 113.49 | 1.04 | 0.73 | -0.32 | 0.75 |

Statistical outputs include the sex effect for the linear mixed model (LMM), the results of the likelihood ratio test, and the results for each pairwise comparison between states of wake (W), anesthesia (A), drug (D), and recovery (R) for saline, DOI, or DOI + volinanserin (DOI + Vol). Corrected p values using Tukey's method are reported for pairwise comparisons. CI = 95% confidence interval; df = degrees of freedom; Estimate = difference in means.

**Table S6. Statistical Outputs for High Gamma Relative Power before and after Infusion of Saline, DOI, or DOI with Volinanserin Pretreatment during Isoflurane Anesthesia**

| | Estimate | df | *t* | *p* | Lower CI | Upper CI |
|---|---|---|---|---|---|---|
| ***LMM: Female v. Male*** | 0.00050 | 9.79 | 1.00 | 0.34 | -0.00044 | 0.0015 |
| | | **df** | $X^2$ | ***p*** | | |
| ***Likelihood Ratio Test*** | - | 6.00 | 26.42 | 0.00019 | - | - |
| | **Estimate** | **df** | ***t*** | ***p*** | **Lower CI** | **Upper CI** |
| ***Saline*** | | | | | | |
| *A ~ W* | -0.0058 | 113.83 | -7.09 | < 0.0001 | -0.0079 | -0.0036 |
| *D ~ W* | -0.0057 | 113.83 | -7.08 | < 0.0001 | -0.0079 | -0.0036 |
| *D ~ A* | 0.000014 | 113.83 | 0.017 | 1.00 | -0.0021 | 0.0021 |
| *R ~ W* | 0.0015 | 114.29 | 1.81 | 0.27 | -0.00067 | 0.0037 |
| *R ~ A* | 0.0073 | 114.29 | 8.72 | < 0.0001 | 0.0051 | 0.0094 |
| *R ~ D* | 0.0073 | 114.29 | 8.70 | < 0.0001 | 0.0051 | 0.0094 |
| ***DOI*** | | | | | | |
| *A ~ W* | -0.0056 | 113.83 | -7.80 | < 0.0001 | -0.0075 | -0.0037 |
| *D ~ W* | -0.0023 | 113.83 | -3.23 | 0.0089 | -0.0042 | -0.00045 |
| *D ~ A* | 0.0033 | 113.83 | 4.57 | < 0.0001 | 0.0014 | 0.0052 |
| *R ~ W* | 0.00089 | 116.90 | 1.15 | 0.66 | -0.0011 | 0.0029 |
| *R ~ A* | 0.0065 | 116.90 | 8.42 | < 0.0001 | 0.0045 | 0.0085 |
| *R ~ D* | 0.0032 | 116.90 | 4.16 | 0.00035 | 0.0012 | 0.0052 |
| ***DOI + Vol*** | | | | | | |
| *A ~ W* | -0.0043 | 113.83 | -5.32 | < 0.0001 | -0.0064 | -0.0022 |
| *D ~ W* | -0.0043 | 113.83 | -5.30 | < 0.0001 | -0.0064 | -0.0022 |
| *D ~ A* | 0.000018 | 113.83 | 0.022 | 1.00 | -0.0021 | 0.0021 |
| *R ~ W* | 0.0037 | 114.66 | 4.35 | 0.00017 | 0.0015 | 0.0060 |
| *R ~ A* | 0.0081 | 114.66 | 9.38 | < 0.0001 | 0.0058 | 0.010 |
| *R ~ D* | 0.0080 | 114.66 | 9.36 | < 0.0001 | 0.0058 | 0.010 |

Statistical outputs include the sex effect for the linear mixed model (LMM), the results of the likelihood ratio test, and the results for each pairwise comparison between states of wake (W), anesthesia (A), drug (D), and recovery (R) for saline, DOI, or DOI + volinanserin (DOI + Vol). Corrected p values using Tukey’s method are reported for pairwise comparisons. CI = 95% confidence interval; df = degrees of freedom; Estimate = difference in means.

**Table S7. Statistical Outputs for Theta Global Coherence before and after Infusion of Saline, DOI, or DOI with Volinanserin Pretreatment during Propofol Anesthesia**

| | **Estimate** | **df** | ***t*** | ***p*** | **Lower CI** | **Upper CI** |
|---|---|---|---|---|---|---|
| ***LMM: Female v. Male*** | -0.055 | 16.28 | -0.45 | 0.66 | -0.30 | 0.18 |
| | | **df** | $X^2$ | ***p*** | | |
| ***Likelihood Ratio Test*** | - | 6.00 | 35.35 | < 0.0001 | - | - |
| | **Estimate** | **df** | ***t*** | ***p*** | **Lower CI** | **Upper CI** |
| ***Saline*** | | | | | | |
| *A ~ W* | -1.17 | 148.11 | -10.14 | < 0.0001 | -1.47 | -0.87 |
| *D ~ W* | -1.11 | 148.11 | -9.60 | < 0.0001 | -1.41 | -0.81 |
| *D ~ A* | 0.062 | 148.11 | 0.54 | 0.95 | -0.24 | 0.36 |
| *R ~ W* | -0.74 | 149.86 | -5.43 | < 0.0001 | -1.09 | -0.39 |
| *R ~ A* | 0.43 | 149.86 | 3.18 | 0.0096 | 0.079 | 0.79 |
| *R ~ D* | 0.37 | 149.86 | 2.72 | 0.036 | 0.016 | 0.72 |
| ***DOI*** | | | | | | |
| *A ~ W* | -1.16 | 148.11 | -10.58 | < 0.0001 | -1.44 | -0.87 |
| *D ~ W* | -1.67 | 148.11 | -15.32 | < 0.0001 | -1.96 | -1.39 |
| *D ~ A* | -0.52 | 148.11 | -4.74 | < 0.0001 | -0.80 | -0.23 |
| *R ~ W* | -0.71 | 149.44 | -5.79 | < 0.0001 | -1.026 | -0.39 |
| *R ~ A* | 0.45 | 149.44 | 3.66 | 0.0020 | 0.13 | 0.77 |
| *R ~ D* | 0.96 | 149.44 | 7.89 | < 0.0001 | 0.65 | 1.28 |
| ***DOI + Vol*** | | | | | | |
| *A ~ W* | -0.92 | 148.11 | -6.73 | < 0.0001 | -1.28 | -0.57 |
| *D ~ W* | -1.06 | 148.11 | -7.74 | < 0.0001 | -1.42 | -0.71 |
| *D ~ A* | -0.14 | 148.11 | -1.01 | 0.74 | -0.50 | 0.22 |
| *R ~ W* | -1.01 | 148.11 | -7.35 | < 0.0001 | -1.37 | -0.65 |
| *R ~ A* | -0.085 | 148.11 | -0.62 | 0.93 | -0.44 | 0.27 |
| *R ~ D* | 0.053 | 148.11 | 0.39 | 0.98 | -0.30 | 0.41 |

Statistical outputs include the sex effect for the linear mixed model (LMM), the results of the likelihood ratio test, and the results for each pairwise comparison between states of wake (W), anesthesia (A), drug (D), and recovery (R) for saline, DOI, or DOI + volinanserin (DOI + Vol). Corrected p values using Tukey's method are reported for pairwise comparisons. CI = 95% confidence interval; df = degrees of freedom; Estimate = difference in means.

**Table S8. Statistical Outputs for High Gamma Global Coherence before and after Infusion of Saline, DOI, or DOI with Volinanserin Pretreatment during Propofol Anesthesia**

| | Estimate | df | *t* | *p* | Lower CI | Upper CI |
|---|---|---|---|---|---|---|
| ***LMM: Female v. Male*** | -0.11 | 16.64 | -0.90 | 0.38 | -0.34 | 0.12 |
| | | **df** | $X^2$ | ***p*** | | |
| ***Likelihood Ratio Test*** | - | 6.00 | 189.75 | < 0.0001 | - | - |
| | **Estimate** | **df** | ***t*** | ***p*** | **Lower CI** | **Upper CI** |
| ***Saline*** | | | | | | |
| *A ~ W* | -1.33 | 148.05 | -14.64 | < 0.0001 | -1.57 | -1.10 |
| *D ~ W* | -1.30 | 148.05 | -14.32 | < 0.0001 | -1.54 | -1.07 |
| *D ~ A* | 0.030 | 148.05 | 0.33 | 0.99 | -0.21 | 0.27 |
| *R ~ W* | -0.48 | 149.24 | -4.45 | < 0.0001 | -0.76 | -0.20 |
| *R ~ A* | 0.86 | 149.24 | 7.98 | < 0.0001 | 0.58 | 1.14 |
| *R ~ D* | 0.83 | 149.24 | 7.70 | < 0.0001 | 0.55 | 1.11 |
| ***DOI*** | | | | | | |
| *A ~ W* | -1.34 | 148.05 | -15.58 | < 0.0001 | -1.57 | -1.12 |
| *D ~ W* | 0.50 | 148.05 | 5.79 | < 0.0001 | 0.28 | 0.72 |
| *D ~ A* | 1.84 | 148.05 | 21.38 | < 0.0001 | 1.62 | 2.07 |
| *R ~ W* | 0.31 | 148.96 | 3.16 | 0.010 | 0.055 | 0.56 |
| *R ~ A* | 1.65 | 148.96 | 17.07 | < 0.0001 | 1.40 | 1.90 |
| *R ~ D* | -0.19 | 148.96 | -2.01 | 0.19 | -0.44 | 0.057 |
| ***DOI + Vol*** | | | | | | |
| *A ~ W* | -1.17 | 148.05 | -10.82 | < 0.0001 | -1.46 | -0.89 |
| *D ~ W* | -1.13 | 148.05 | -10.45 | < 0.0001 | -1.42 | -0.85 |
| *D ~ A* | 0.040 | 148.05 | 0.37 | 0.98 | -0.24 | 0.32 |
| *R ~ W* | -0.23 | 148.05 | -2.10 | 0.16 | -0.51 | 0.054 |
| *R ~ A* | 0.95 | 148.05 | 8.72 | < 0.0001 | 0.66 | 1.23 |
| *R ~ D* | 0.91 | 148.05 | 8.35 | < 0.0001 | 0.62 | 1.19 |

Statistical outputs include the sex effect for the linear mixed model (LMM), the results of the likelihood ratio test, and the results for each pairwise comparison between states of wake (W), anesthesia (A), drug (D), and recovery (R) for saline, DOI, or DOI + volinanserin (DOI + Vol). Corrected p values using Tukey's method are reported for pairwise comparisons. CI = 95% confidence interval; df = degrees of freedom; Estimate = difference in means.

**Table S9. Statistical Outputs for Theta Global Coherence before and after Infusion of Saline, DOI, or DOI with Volinanserin Pretreatment during Isoflurane Anesthesia**

| | **Estimate** | **df** | ***t*** | ***p*** | **Lower CI** | **Upper CI** |
|---|---|---|---|---|---|---|
| ***LMM: Female v. Male*** | -0.16 | 10.86 | -1.09 | 0.30 | -0.44 | 0.12 |
| | | **df** | ***$X^2$*** | ***p*** | | |
| ***Likelihood Ratio Test*** | - | 6.00 | 48.16 | < 0.0001 | - | - |
| | **Estimate** | **df** | ***t*** | ***p*** | **Lower CI** | **Upper CI** |
| ***Saline*** | | | | | | |
| *A ~ W* | -0.74 | 113.15 | -5.89 | < 0.0001 | -1.06 | -0.41 |
| *D ~ W* | -0.75 | 113.15 | -6.00 | < 0.0001 | -1.08 | -0.43 |
| *D ~ A* | -0.014 | 113.15 | -0.11 | 1.00 | -0.34 | 0.31 |
| *R ~ W* | -0.95 | 113.28 | -7.35 | < 0.0001 | -1.28 | -0.61 |
| *R ~ A* | -0.21 | 113.28 | -1.62 | 0.37 | -0.54 | 0.13 |
| *R ~ D* | -0.19 | 113.28 | -1.51 | 0.44 | -0.53 | 0.14 |
| ***DOI*** | | | | | | |
| *A ~ W* | -0.85 | 113.15 | -7.65 | < 0.0001 | -1.14 | -0.56 |
| *D ~ W* | -1.76 | 113.15 | -15.89 | < 0.0001 | -2.05 | -1.48 |
| *D ~ A* | -0.92 | 113.15 | -8.24 | < 0.0001 | -1.20 | -0.63 |
| *R ~ W* | -1.18 | 115.21 | -9.83 | < 0.0001 | -1.49 | -0.87 |
| *R ~ A* | -0.33 | 115.21 | -2.76 | 0.034 | -0.64 | -0.018 |
| *R ~ D* | 0.58 | 115.21 | 4.85 | < 0.0001 | 0.27 | 0.90 |
| ***DOI + Vol*** | | | | | | |
| *~ W* | -1.07 | 113.15 | -8.57 | < 0.0001 | -1.40 | -0.75 |
| *D ~ W* | -1.15 | 113.15 | -9.21 | < 0.0001 | -1.48 | -0.83 |
| *D ~ A* | -0.080 | 113.15 | -0.64 | 0.92 | -0.41 | 0.25 |
| *R ~ W* | -1.09 | 113.38 | -8.18 | < 0.0001 | -1.43 | -0.74 |
| *R ~ A* | -0.010 | 113.38 | -0.078 | 1.00 | -0.36 | 0.34 |
| *R ~ D* | 0.070 | 113.38 | 0.53 | 0.95 | -0.28 | 0.42 |

Statistical outputs include the sex effect for the linear mixed model (LMM), the results of the likelihood ratio test, and the results for each pairwise comparison between states of wake (W), anesthesia (A), drug (D), and recovery (R) for saline, DOI, or DOI + volinanserin (DOI + Vol). Corrected p values using Tukey’s method are reported for pairwise comparisons. CI = 95% confidence interval; df = degrees of freedom; Estimate = difference in means.

**Table S10. Statistical Outputs for High Gamma Global Coherence before and after Infusion of Saline, DOI, or DOI with Volinanserin Pretreatment during Isoflurane Anesthesia**

| | Estimate | df | *t* | *p* | Lower CI | Upper CI |
|---|---|---|---|---|---|---|
| ***LMM: Female v. Male*** | 0.063 | 10.04 | 0.53 | 0.61 | -0.18 | 0.29 |
| | | **df** | $X^2$ | ***p*** | | |
| ***Likelihood Ratio Test*** | - | 6.00 | 107.60 | < 0.0001 | - | - |
| | **Estimate** | **df** | ***t*** | ***p*** | **Lower CI** | **Upper CI** |
| ***Saline*** | | | | | | |
| *A ~ W* | -1.44 | 113.65 | -8.25 | < 0.0001 | -1.89 | -0.98 |
| *D ~ W* | -1.42 | 113.65 | -8.14 | < 0.0001 | -1.87 | -0.96 |
| *D ~ A* | 0.018 | 113.65 | 0.11 | 1.00 | -0.44 | 0.47 |
| *R ~ W* | -0.20 | 114.039 | -1.14 | 0.67 | -0.67 | 0.26 |
| *R ~ A* | 1.23 | 114.039 | 6.90 | < 0.0001 | 0.77 | 1.70 |
| *R ~ D* | 1.21 | 114.039 | 6.79 | < 0.0001 | 0.75 | 1.68 |
| ***DOI*** | | | | | | |
| *A ~ W* | -1.21 | 113.65 | -7.84 | < 0.0001 | -1.61 | -0.81 |
| *D ~ W* | 0.81 | 113.65 | 5.25 | < 0.0001 | 0.41 | 1.21 |
| *D ~ A* | 2.020 | 113.65 | 13.090 | < 0.0001 | 1.62 | 2.42 |
| *R ~ W* | 0.37 | 116.70 | 2.25 | 0.12 | -0.059 | 0.81 |
| *R ~ A* | 1.58 | 116.70 | 9.54 | < 0.0001 | 1.15 | 2.017 |
| *R ~ D* | -0.44 | 116.70 | -2.62 | 0.048 | -0.87 | -0.0030 |
| ***DOI + Vol*** | | | | | | |
| *A ~ W* | -1.60 | 113.65 | -9.17 | < 0.0001 | -2.050 | -1.14 |
| *D ~ W* | -1.48 | 113.65 | -8.50 | < 0.0001 | -1.93 | -1.026 |
| *D ~ A* | 0.12 | 113.65 | 0.67 | 0.91 | -0.34 | 0.57 |
| *R ~ W* | -0.24 | 114.33 | -1.33 | 0.55 | -0.73 | 0.24 |
| *R ~ A* | 1.35 | 114.33 | 7.34 | < 0.0001 | 0.87 | 1.83 |
| *R ~ D* | 1.24 | 114.33 | 6.71 | < 0.0001 | 0.76 | 1.72 |

Statistical outputs include the sex effect for the linear mixed model (LMM), the results of the likelihood ratio test, and the results for each pairwise comparison between states of wake (W), anesthesia (A), drug (D), and recovery (R) for saline, DOI, or DOI + volinanserin (DOI + Vol). Corrected p values using Tukey's method are reported for pairwise comparisons. CI = 95% confidence interval; df = degrees of freedom; Estimate = difference in means.

**Table S11. Statistical Outputs for Theta Feedforward Connectivity before and after Infusion of Saline, DOI, or DOI with Volinanserin Pretreatment during Propofol Anesthesia**

| | Estimate | df | *t* | *p* | Lower CI | Upper CI |
|---|---|---|---|---|---|---|
| ***LMM: Female v. Male*** | -0.00073 | 17.15 | -0.20 | 0.84 | -0.0079 | 0.0064 |
| | | **df** | $X^2$ | ***p*** | | |
| ***Likelihood Ratio Test*** | - | 6.00 | 43.85 | < 0.0001 | - | - |
| | **Estimate** | **df** | ***t*** | ***p*** | **Lower CI** | **Upper CI** |
| ***Saline*** | | | | | | |
| *A ~ W* | -0.032 | 148.14 | -8.65 | < 0.0001 | -0.042 | -0.022 |
| *D ~ W* | -0.029 | 148.14 | -7.82 | < 0.0001 | -0.039 | -0.019 |
| *D ~ A* | 0.0030 | 148.14 | 0.82 | 0.84 | -0.0066 | 0.013 |
| *R ~ W* | -0.023 | 150.13 | -5.21 | < 0.0001 | -0.034 | -0.011 |
| *R ~ A* | 0.0093 | 150.13 | 2.14 | 0.15 | -0.0020 | 0.021 |
| *R ~ D* | 0.0062 | 150.13 | 1.44 | 0.48 | -0.0051 | 0.018 |
| ***DOI*** | | | | | | |
| *A ~ W* | -0.038 | 148.14 | -10.72 | < 0.0001 | -0.047 | -0.028 |
| *D ~ W* | -0.054 | 148.14 | -15.31 | < 0.0001 | -0.063 | -0.044 |
| *D ~ A* | -0.016 | 148.14 | -4.58 | < 0.0001 | -0.025 | -0.0069 |
| *R ~ W* | -0.015 | 149.65 | -3.89 | 0.00086 | -0.025 | -0.0051 |
| *R ~ A* | 0.022 | 149.65 | 5.68 | < 0.0001 | 0.012 | 0.032 |
| *R ~ D* | 0.038 | 149.65 | 9.78 | < 0.0001 | 0.028 | 0.048 |
| ***DOI + Vol*** | | | | | | |
| *A ~ W* | -0.030 | 148.14 | -6.93 | < 0.0001 | -0.042 | -0.019 |
| *D ~ W* | -0.031 | 148.14 | -7.13 | < 0.0001 | -0.043 | -0.020 |
| *D ~ A* | -0.0009 | 148.14 | -0.21 | 1.00 | -0.012 | 0.011 |
| *R ~ W* | -0.023 | 148.14 | -5.13 | < 0.0001 | -0.034 | -0.011 |
| *R ~ A* | 0.0079 | 148.14 | 1.79 | 0.28 | -0.0035 | 0.019 |
| *R ~ D* | 0.0088 | 148.14 | 2.00 | 0.19 | -0.0026 | 0.020 |

Statistical outputs include the sex effect for the linear mixed model (LMM), the results of the likelihood ratio test, and the results for each pairwise comparison between states of wake (W), anesthesia (A), drug (D), and recovery (R) for saline, DOI, or DOI + volinanserin (DOI + Vol). Corrected p values using Tukey's method are reported for pairwise comparisons. CI = 95% confidence interval; df = degrees of freedom; Estimate = difference in means.

**Table S12. Statistical Outputs for High Gamma Feedforward Connectivity before and after Infusion of Saline, DOI, or DOI with Volinanserin Pretreatment during Propofol Anesthesia**

| | Estimate | df | *t* | *p* | Lower CI | Upper CI |
|---|---|---|---|---|---|---|
| ***LMM: Female v. Male*** | 0.00018 | 18.21 | 0.15 | 0.88 | -0.0022 | 0.0026 |
| | | **df** | **$X^2$** | ***p*** | | |
| ***Likelihood Ratio Test*** | - | 6.00 | 143.30 | < 0.0001 | - | - |
| | **Estimate** | **df** | ***t*** | ***p*** | **Lower CI** | **Upper CI** |
| ***Saline*** | | | | | | |
| *A ~ W* | -0.014 | 148.09 | -12.55 | < 0.0001 | -0.017 | -0.011 |
| *D ~ W* | -0.013 | 148.09 | -11.93 | < 0.0001 | -0.016 | -0.010 |
| *D ~ A* | 0.00069 | 148.09 | 0.62 | 0.93 | -0.0022 | 0.0036 |
| *R ~ W* | -0.0063 | 149.70 | -4.82 | < 0.0001 | -0.0098 | -0.0029 |
| *R ~ A* | 0.0077 | 149.70 | 5.84 | < 0.0001 | 0.0043 | 0.011 |
| *R ~ D* | 0.0070 | 149.70 | 5.31 | < 0.0001 | 0.0036 | 0.010 |
| ***DOI*** | | | | | | |
| *A ~ W* | -0.013 | 148.09 | -12.65 | < 0.0001 | -0.016 | -0.011 |
| *D ~ W* | 0.0038 | 148.09 | 3.58 | 0.0026 | 0.0010 | 0.0065 |
| *D ~ A* | 0.017 | 148.09 | 16.23 | < 0.0001 | 0.014 | 0.020 |
| *R ~ W* | 0.0048 | 149.32 | 4.03 | 0.00051 | 0.0017 | 0.0079 |
| *R ~ A* | 0.018 | 149.32 | 15.32 | < 0.0001 | 0.015 | 0.021 |
| *R ~ D* | 0.0010 | 149.32 | 0.83 | 0.84 | -0.0021 | 0.0041 |
| ***DOI + Vol*** | | | | | | |
| *A ~ W* | -0.012 | 148.09 | -8.99 | < 0.0001 | -0.015 | -0.0085 |
| *D ~ W* | -0.012 | 148.09 | -9.10 | < 0.0001 | -0.016 | -0.0086 |
| *D ~ A* | -0.00015 | 148.09 | -0.11 | 1.00 | -0.0036 | 0.0033 |
| *R ~ W* | -0.0039 | 148.09 | -2.90 | 0.022 | -0.0073 | -0.00041 |
| *R ~ A* | 0.0081 | 148.09 | 6.08 | < 0.0001 | 0.0046 | 0.012 |
| *R ~ D* | 0.0082 | 148.09 | 6.19 | < 0.0001 | 0.0048 | 0.012 |

Statistical outputs include the sex effect for the linear mixed model (LMM), the results of the likelihood ratio test, and the results for each pairwise comparison between states of wake (W), anesthesia (A), drug (D), and recovery (R) for saline, DOI, or DOI + volinanserin (DOI + Vol). Corrected p values using Tukey's method are reported for pairwise comparisons. CI = 95% confidence interval; df = degrees of freedom; Estimate = difference in means.

**Table S13. Statistical Outputs for Theta Feedforward Connectivity before and after Infusion of Saline, DOI, or DOI with Volinanserin Pretreatment during Isoflurane Anesthesia**

| | **Estimate** | **df** | ***t*** | ***p*** | **Lower CI** | **Upper CI** |
|---|---|---|---|---|---|---|
| ***LMM: Female v. Male*** | 0.013 | 10.80 | -2.80 | 0.018 | -0.022 | -0.0040 |
| | | **df** | **$X^2$** | ***p*** | | |
| ***Likelihood Ratio Test*** | - | 6.00 | 8.075 | 0.23 | - | - |
| | **Estimate** | **df** | ***t*** | ***p*** | **Lower CI** | **Upper CI** |
| ***Saline*** | | | | | | |
| *A ~ W* | -0.046 | 113.34 | -8.84 | < 0.0001 | -0.059 | -0.032 |
| *D ~ W* | -0.046 | 113.34 | -8.97 | < 0.0001 | -0.060 | -0.033 |
| *D ~ A* | -0.00069 | 113.34 | -0.13 | 1.00 | -0.014 | 0.013 |
| *R ~ W* | -0.031 | 113.57 | -5.86 | < 0.0001 | -0.045 | -0.017 |
| *R ~ A* | 0.015 | 113.57 | 2.75 | 0.034 | 0.00076 | 0.028 |
| *R ~ D* | 0.015 | 113.57 | 2.88 | 0.024 | 0.0014 | 0.029 |
| ***DOI*** | | | | | | |
| *A ~ W* | -0.046 | 113.34 | -10.058 | < 0.0001 | -0.058 | -0.034 |
| *D ~ W* | -0.060 | 113.34 | -13.0094 | < 0.0001 | -0.071 | -0.048 |
| *D ~ A* | -0.014 | 113.34 | -2.95 | 0.020 | -0.025 | -0.0016 |
| *R ~ W* | -0.034 | 116.03 | -6.97 | < 0.0001 | -0.047 | -0.022 |
| *R ~ A* | 0.012 | 116.03 | 2.35 | 0.093 | -0.0013 | 0.024 |
| *R ~ D* | 0.025 | 116.03 | 5.082 | < 0.0001 | 0.012 | 0.038 |
| ***DOI + Vol*** | | | | | | |
| *A ~ W* | -0.056 | 113.34 | -10.78 | < 0.0001 | -0.069 | -0.042 |
| *D ~ W* | -0.056 | 113.34 | -10.91 | < 0.0001 | -0.070 | -0.043 |
| *D ~ A* | -0.00068 | 113.34 | -0.13 | 1.00 | -0.014 | 0.013 |
| *R ~ W* | -0.035 | 113.74 | -6.40 | < 0.0001 | -0.049 | -0.021 |
| *R ~ A* | 0.021 | 113.74 | 3.79 | 0.0014 | 0.0065 | 0.035 |
| *R ~ D* | 0.021 | 113.74 | 3.91 | 0.00088 | 0.0071 | 0.036 |

Statistical outputs include the sex effect for the linear mixed model (LMM), the results of the likelihood ratio test, and the results for each pairwise comparison between states of wake (W), anesthesia (A), drug (D), and recovery (R) for saline, DOI, or DOI + volinanserin (DOI + Vol). Corrected p values using Tukey’s method are reported for pairwise comparisons. CI = 95% confidence interval; df = degrees of freedom; Estimate = difference in means.

**Table S14. Statistical Outputs for High Gamma Feedforward Connectivity before and after Infusion of Saline, DOI, or DOI with Volinanserin Pretreatment during Isoflurane Anesthesia**

| | **Estimate** | **df** | ***t*** | ***p*** | **Lower CI** | **Upper CI** |
|---|---|---|---|---|---|---|
| ***LMM: Female v. Male*** | 0.0015 | 10.78 | 1.15 | 0.28 | -0.0011 | 0.0041 |
| | | **df** | **$X^2$** | ***p*** | | |
| ***Likelihood Ratio Test*** | - | 6.00 | 88.79 | < 0.0001 | - | - |
| | **Estimate** | **df** | ***t*** | ***p*** | **Lower CI** | **Upper CI** |
| ***Saline*** | | | | | | |
| *A ~ W* | -0.012 | 113.55 | -6.58 | < 0.0001 | -0.017 | -0.0074 |
| *D ~ W* | -0.013 | 113.55 | -6.84 | < 0.0001 | -0.018 | -0.0078 |
| *D ~ A* | -0.00048 | 113.55 | -0.26 | 0.99 | -0.0053 | 0.0044 |
| *R ~ W* | -0.0047 | 113.89 | -2.49 | 0.067 | -0.0097 | 0.00023 |
| *R ~ A* | 0.0075 | 113.89 | 3.92 | 0.00086 | 0.0025 | 0.012 |
| *R ~ D* | 0.0079 | 113.89 | 4.17 | 0.00034 | 0.0030 | 0.013 |
| ***DOI*** | | | | | | |
| *A ~ W* | -0.0096 | 113.55 | -5.83 | < 0.0001 | -0.014 | -0.0053 |
| *D ~ W* | 0.0066 | 113.55 | 4.03 | 0.00058 | 0.0023 | 0.011 |
| *D ~ A* | 0.016 | 113.55 | 9.86 | < 0.0001 | 0.012 | 0.021 |
| *R ~ W* | -0.0031 | 116.54 | -1.77 | 0.29 | -0.0077 | 0.0015 |
| *R ~ A* | 0.0064 | 116.54 | 3.64 | < 0.0001 | 0.0018 | 0.011 |
| *R ~ D* | -0.0097 | 116.54 | -5.51 | < 0.0001 | -0.014 | -0.0051 |
| ***DOI + Vol*** | | | | | | |
| *A ~ W* | -0.016 | 113.55 | -8.40 | < 0.0001 | -0.020 | -0.011 |
| *D ~ W* | -0.015 | 113.55 | -8.20 | < 0.0001 | -0.0200 | -0.010 |
| *D ~ A* | 0.00036 | 113.55 | 0.19 | 1.00 | -0.0045 | 0.0052 |
| *R ~ W* | -0.0061 | 114.14 | -3.12 | 0.012 | -0.011 | -0.0010 |
| *R ~ A* | 0.0094 | 114.14 | 4.82 | < 0.0001 | 0.0043 | 0.015 |
| *R ~ D* | 0.0091 | 114.14 | 4.63 | < 0.0001 | 0.0040 | 0.014 |

Statistical outputs include the sex effect for the linear mixed model (LMM), the results of the likelihood ratio test, and the results for each pairwise comparison between states of wake (W), anesthesia (A), drug (D), and recovery (R) for saline, DOI, or DOI + volinanserin (DOI + Vol). Corrected p values using Tukey’s method are reported for pairwise comparisons. CI = 95% confidence interval; df = degrees of freedom; Estimate = difference in means.

**Table S15. Statistical Outputs for Theta Feedback Connectivity before and after Infusion of Saline, DOI, or DOI with Volinanserin Pretreatment during Propofol Anesthesia**

| | Estimate | df | *t* | *p* | Lower CI | Upper CI |
|---|---|---|---|---|---|---|
| ***LMM: Female v. Male*** | -0.0026 | 16.94 | -0.94 | 0.36 | -0.0081 | 0.0028 |
| | | **df** | $X^2$ | ***p*** | | |
| ***Likelihood Ratio Test*** | - | 6.00 | 19.27 | 0.0037 | - | - |
| | **Estimate** | **df** | ***t*** | ***p*** | **Lower CI** | **Upper CI** |
| ***Saline*** | | | | | | |
| *A ~ W* | -0.024 | 148.34 | -6.51 | < 0.0001 | -0.033 | -0.014 |
| *D ~ W* | -0.024 | 148.34 | -6.43 | < 0.0001 | -0.033 | -0.014 |
| *D ~ A* | 0.00029 | 148.34 | 0.079 | 1.00 | -0.0092 | 0.0098 |
| *R ~ W* | -0.011 | 151.46 | -2.59 | 0.051 | -0.022 | 0.000036 |
| *R ~ A* | 0.013 | 151.46 | 2.95 | 0.019 | 0.0015 | 0.024 |
| *R ~ D* | 0.012 | 151.46 | 2.89 | 0.023 | 0.0012 | 0.024 |
| ***DOI*** | | | | | | |
| *A ~ W* | -0.032 | 148.34 | -9.37 | < 0.0001 | -0.041 | -0.023 |
| *D ~ W* | -0.039 | 148.34 | -11.17 | < 0.0001 | -0.048 | -0.030 |
| *D ~ A* | -0.0062 | 148.34 | -1.80 | 0.28 | -0.015 | 0.0028 |
| *R ~ W* | -0.014 | 150.69 | -3.73 | 0.0015 | -0.025 | -0.0044 |
| *R ~ A* | 0.018 | 150.69 | 4.64 | < 0.0001 | 0.0079 | 0.028 |
| *R ~ D* | 0.024 | 150.69 | 6.25 | < 0.0001 | 0.014 | 0.034 |
| ***DOI + Vol*** | | | | | | |
| *A ~ W* | -0.021 | 148.34 | -4.86 | < 0.0001 | -0.032 | -0.0099 |
| *D ~ W* | -0.023 | 148.34 | -5.35 | < 0.0001 | -0.035 | -0.012 |
| *D ~ A* | -0.0021 | 148.34 | -0.49 | 0.96 | -0.013 | 0.0092 |
| *R ~ W* | -0.018 | 148.34 | -4.18 | 0.00029 | -0.030 | -0.0069 |
| *R ~ A* | 0.0030 | 148.34 | 0.69 | 0.90 | -0.0083 | 0.014 |
| *R ~ D* | 0.0051 | 148.34 | 1.17 | 0.65 | -0.0062 | 0.016 |

Statistical outputs include the sex effect for the linear mixed model (LMM), the results of the likelihood ratio test, and the results for each pairwise comparison between states of wake (W), anesthesia (A), drug (D), and recovery (R) for saline, DOI, or DOI + volinanserin (DOI + Vol). Corrected p values using Tukey's method are reported for pairwise comparisons. CI = 95% confidence interval; df = degrees of freedom; Estimate = difference in means.

**Table S16. Statistical Outputs for High Gamma Feedback Connectivity before and after Infusion of Saline, DOI, or DOI with Volinanserin Pretreatment during Propofol Anesthesia**

| | Estimate | df | *t* | *p* | Lower CI | Upper CI |
|---|---|---|---|---|---|---|
| ***LMM: Female v. Male*** | 0.0018 | 19.09 | 1.23 | 0.24 | -0.0011 | 0.0047 |
| | | **df** | $X^2$ | ***p*** | | |
| ***Likelihood Ratio Test*** | - | 6.00 | 103.88 | < 0.0001 | - | - |
| | **Estimate** | **df** | ***t*** | ***p*** | **Lower CI** | **Upper CI** |
| ***Saline*** | | | | | | |
| *A ~ W* | -0.014 | 148.09 | -10.99 | < 0.0001 | -0.018 | -0.011 |
| *D ~ W* | -0.014 | 148.09 | -10.54 | < 0.0001 | -0.017 | -0.010 |
| *D ~ A* | 0.00058 | 148.09 | 0.45 | 0.97 | -0.0028 | 0.0040 |
| *R ~ W* | -0.0074 | 149.61 | -4.85 | < 0.0001 | -0.011 | -0.0035 |
| *R ~ A* | 0.0069 | 149.61 | 4.49 | < 0.0001 | 0.0029 | 0.011 |
| *R ~ D* | 0.0063 | 149.61 | 4.11 | 0.00038 | 0.0023 | 0.010 |
| ***DOI*** | | | | | | |
| *A ~ W* | -0.015 | 148.09 | -11.73 | < 0.0001 | -0.018 | -0.011 |
| *D ~ W* | 0.0019 | 148.09 | 1.54 | 0.42 | -0.0013 | 0.0051 |
| *D ~ A* | 0.016 | 148.09 | 13.27 | < 0.0001 | 0.013 | 0.020 |
| *R ~ W* | 0.0029 | 149.25 | 2.13 | 0.15 | -0.00065 | 0.0065 |
| *R ~ A* | 0.017 | 149.25 | 12.60 | < 0.0001 | 0.014 | 0.021 |
| *R ~ D* | 0.0010 | 149.25 | 0.75 | 8.74E-01 | -0.0025 | 0.0046 |
| ***DOI + Vol*** | | | | | | |
| *A ~ W* | -0.012 | 148.09 | -7.81 | < 0.0001 | -0.016 | -0.0081 |
| *D ~ W* | -0.012 | 148.09 | -7.61 | < 0.0001 | -0.016 | -0.0078 |
| *D ~ A* | 0.00032 | 148.09 | 0.20 | 1.00 | -0.0037 | 0.0043 |
| *R ~ W* | -0.0040 | 148.09 | -2.60 | 0.049 | -0.0081 | -0.0000079 |
| *R ~ A* | 0.0081 | 148.09 | 5.21 | < 0.0001 | 0.0041 | 0.012 |
| *R ~ D* | 0.0078 | 148.09 | 5.01 | < 0.0001 | 0.0037 | 0.012 |

Statistical outputs include the sex effect for the linear mixed model (LMM), the results of the likelihood ratio test, and the results for each pairwise comparison between states of wake (W), anesthesia (A), drug (D), and recovery (R) for saline, DOI, or DOI + volinanserin (DOI + Vol). Corrected p values using Tukey's method are reported for pairwise comparisons. CI = 95% confidence interval; df = degrees of freedom; Estimate = difference in means.

**Table S17. Statistical Outputs for Theta Feedback Connectivity before and after Infusion of Saline, DOI, or DOI with Volinanserin Pretreatment during Isoflurane Anesthesia**

| | Estimate | df | *t* | *p* | Lower CI | Upper CI |
|---|---|---|---|---|---|---|
| ***LMM: Female v. Male*** | -0.011 | 9.97 | -2.54 | 0.029 | -0.020 | -0.0028 |
| | | **df** | $X^2$ | ***p*** | | |
| ***Likelihood Ratio Test*** | - | 6.00 | 9.89 | 0.13 | - | - |
| | **Estimate** | **df** | ***t*** | ***p*** | **Lower CI** | **Upper CI** |
| ***Saline*** | | | | | | |
| *A ~ W* | -0.030 | 113.24 | -6.65 | < 0.0001 | -0.042 | -0.018 |
| *D ~ W* | -0.032 | 113.24 | -7.11 | < 0.0001 | -0.044 | -0.020 |
| *D ~ A* | -0.0021 | 113.24 | -0.45 | 0.97 | -0.014 | 0.010 |
| *R ~ W* | -0.023 | 113.42 | -4.91 | < 0.0001 | -0.035 | -0.011 |
| *R ~ A* | 0.0073 | 113.42 | 1.57 | 0.40 | -0.0048 | 0.019 |
| *R ~ D* | 0.0094 | 113.42 | 2.01 | 0.19 | -0.0028 | 0.022 |
| ***DOI*** | | | | | | |
| *A ~ W* | -0.041 | 113.24 | -10.20 | < 0.0001 | -0.052 | -0.031 |
| *D ~ W* | -0.048 | 113.24 | -11.87 | < 0.0001 | -0.058 | -0.037 |
| *D ~ A* | -0.0067 | 113.24 | -1.68 | 0.34 | -0.017 | 0.0038 |
| *R ~ W* | -0.030 | 115.67 | -6.89 | < 0.0001 | -0.041 | -0.019 |
| *R ~ A* | 0.011 | 115.67 | 2.54 | 0.059 | 0.00028 | 0.022 |
| *R ~ D* | 0.018 | 115.67 | 4.09 | 0.00046 | 0.0065 | 0.029 |
| ***DOI + Vol*** | | | | | | |
| *A ~ W* | -0.044 | 113.24 | -9.59 | < 0.0001 | -0.055 | -0.032 |
| *D ~ W* | -0.045 | 113.24 | -9.79 | < 0.0001 | -0.056 | -0.033 |
| *D ~ A* | -0.00095 | 113.24 | -0.21 | 1.00 | -0.013 | 0.011 |
| *R ~ W* | -0.027 | 113.55 | -5.53 | < 0.0001 | -0.039 | -0.014 |
| *R ~ A* | 0.017 | 113.55 | 3.53 | 0.0034 | 0.0044 | 0.030 |
| *R ~ D* | 0.018 | 113.55 | 3.72 | 0.0017 | 0.0054 | 0.030 |

Statistical outputs include the sex effect for the linear mixed model (LMM), the results of the likelihood ratio test, and the results for each pairwise comparison between states of wake (W), anesthesia (A), drug (D), and recovery (R) for saline, DOI, or DOI + volinanserin (DOI + Vol). Corrected p values using Tukey's method are reported for pairwise comparisons. CI = 95% confidence interval; df = degrees of freedom; Estimate = difference in means.

**Table S18. Statistical Outputs for High Gamma Feedback Connectivity before and after Infusion of Saline, DOI, or DOI with Volinanserin Pretreatment during Isoflurane Anesthesia**

| | Estimate | df | *t* | *p* | Lower CI | Upper CI |
|---|---|---|---|---|---|---|
| ***LMM: Female v. Male*** | 0.0023 | 11.59 | 1.78 | 0.10 | -.00020 | 0.0047 |
| | | **df** | $X^2$ | ***p*** | | |
| ***Likelihood Ratio Test*** | - | 6.00 | 76.54 | < 0.0001 | - | - |
| | **Estimate** | **df** | ***t*** | ***p*** | **Lower CI** | **Upper CI** |
| ***Saline*** | | | | | | |
| *A ~ W* | -0.012 | 113.64 | -6.20 | < 0.0001 | -0.016 | -0.0067 |
| *D ~ W* | -0.011 | 113.64 | -6.12 | < 0.0001 | -0.016 | -0.0065 |
| *D ~ A* | 0.00015 | 113.64 | 0.081 | 1.00 | -0.0047 | 0.0050 |
| *R ~ W* | -0.0051 | 114.02 | -2.66 | 0.044 | -0.010 | -0.000093 |
| *R ~ A* | 0.0064 | 114.02 | 3.38 | 0.0054 | 0.0015 | 0.011 |
| *R ~ D* | 0.0063 | 114.02 | 3.30 | 0.0069 | 0.0013 | 0.011 |
| ***DOI*** | | | | | | |
| *A ~ W* | -0.0085 | 113.64 | -5.20 | < 0.0001 | -0.013 | -0.0043 |
| *D ~ W* | 0.0054 | 113.64 | 3.26 | 0.0079 | 0.0011 | 0.0097 |
| *D ~ A* | 0.014 | 113.64 | 8.46 | < 0.0001 | 0.010 | 0.018 |
| *R ~ W* | -0.0060 | 116.68 | -3.40 | 0.0051 | -0.011 | -0.0014 |
| *R ~ A* | 0.0025 | 116.68 | 1.43 | 0.048 | -0.0021 | 0.0071 |
| *R ~ D* | -0.011 | 116.68 | -6.43 | < 0.0001 | -0.016 | -0.0068 |
| ***DOI + Vol*** | | | | | | |
| *A ~ W* | -0.014 | 113.64 | -7.47 | < 0.0001 | -0.019 | -0.0090 |
| *D ~ W* | -0.014 | 113.64 | -7.26 | < 0.0001 | -0.018 | -0.0086 |
| *D ~ A* | 0.00039 | 113.64 | 0.21 | 1.00 | -0.0045 | 0.0052 |
| *R ~ W* | -0.0063 | 114.31 | -3.22 | 0.0089 | -0.011 | -0.0012 |
| *R ~ A* | 0.0075 | 114.31 | 3.83 | 0.0012 | 0.0024 | 0.013 |
| *R ~ D* | 0.0071 | 114.31 | 3.64 | 0.0023 | 0.0020 | 0.012 |

Statistical outputs include the sex effect for the linear mixed model (LMM), the results of the likelihood ratio test, and the results for each pairwise comparison between states of wake (W), anesthesia (A), drug (D), and recovery (R) for saline, DOI, or DOI + volinanserin (DOI + Vol). Corrected p values using Tukey's method are reported for pairwise comparisons. CI = 95% confidence interval; df = degrees of freedom; Estimate = difference in means.

**Table S19 Statistical Outputs for Theta Average Node Degree before and after Infusion of Saline, DOI, or DOI with Volinanserin Pretreatment during Propofol Anesthesia**

| | Estimate | df | *t* | *p* | Lower CI | Upper CI |
|---|---|---|---|---|---|---|
| ***LMM: Female v. Male*** | -0.18 | 16.96 | -0.16 | 0.88 | -2.45 | 2.05 |
| | | **df** | $X^2$ | ***p*** | | |
| ***Likelihood Ratio Test*** | - | 6.00 | 36.10 | < 0.0001 | - | - |
| | **Estimate** | **df** | ***t*** | ***p*** | **Lower CI** | **Upper CI** |
| ***Saline*** | | | | | | |
| *A ~ W* | -9.18 | 148.15 | -7.81 | < 0.0001 | -12.23 | -6.12 |
| *D ~ W* | -8.42 | 148.15 | -7.16 | < 0.0001 | -11.47 | -5.36 |
| *D ~ A* | 0.76 | 148.15 | 0.65 | 0.92 | -2.29 | 3.82 |
| *R ~ W* | -4.99 | 150.16 | -3.60 | 0.0024 | -8.58 | -1.39 |
| *R ~ A* | 4.19 | 150.16 | 3.031 | 0.015 | 0.60 | 7.79 |
| *R ~ D* | 3.43 | 150.16 | 2.48 | 0.067 | -0.17 | 7.02 |
| ***DOI*** | | | | | | |
| *A ~ W* | -9.05 | 148.15 | -8.14 | < 0.0001 | -11.94 | -6.16 |
| *D ~ W* | -14.15 | 148.15 | -12.72 | < 0.0001 | -17.04 | -11.26 |
| *D ~ A* | -5.10 | 148.15 | -4.58 | < 0.0001 | -7.99 | -2.21 |
| *R ~ W* | -4.59 | 149.68 | -3.69 | 0.0018 | -7.83 | -1.36 |
| *R ~ A* | 4.46 | 149.68 | 3.58 | 0.0026 | 1.22 | 7.69 |
| *R ~ D* | 9.56 | 149.68 | 7.67 | < 0.0001 | 6.32 | 12.79 |
| ***DOI + Vol*** | | | | | | |
| *A ~ W* | -6.77 | 148.15 | -4.84 | < 0.0001 | -10.40 | -3.13 |
| *D ~ W* | -8.23 | 148.15 | -5.88 | < 0.0001 | -11.87 | -4.60 |
| *D ~ A* | -1.46 | 148.15 | -1.05 | 0.72 | -5.10 | 2.17 |
| *R ~ W* | -8.12 | 148.15 | -5.80 | < 0.0001 | -11.75 | -4.48 |
| *R ~ A* | -1.35 | 148.15 | -0.96 | 0.77 | -4.98 | 2.29 |
| *R ~ D* | 0.12 | 148.15 | 0.083 | 1.00 | -3.52 | 3.75 |

Statistical outputs include the sex effect for the linear mixed model (LMM), the results of the likelihood ratio test, and the results for each pairwise comparison between states of wake (W), anesthesia (A), drug (D), and recovery (R) for saline, DOI, or DOI + volinanserin (DOI + Vol). Corrected p values using Tukey's method are reported for pairwise comparisons. CI = 95% confidence interval; df = degrees of freedom; Estimate = difference in means.

**Table S20 Statistical Outputs for High Gamma Average Node Degree before and after Infusion of Saline, DOI, or DOI with Volinanserin Pretreatment during Propofol Anesthesia**

| | Estimate | df | *t* | *p* | Lower CI | Upper CI |
|---|---|---|---|---|---|---|
| ***LMM: Female v. Male*** | -1.45 | 16.46 | -1.44 | 0.17 | -3.44 | 0.50 |
| | | **df** | $X^2$ | ***p*** | | |
| ***Likelihood Ratio Test*** | - | 6.00 | 214.43 | < 0.0001 | - | - |
| | **Estimate** | **df** | ***t*** | ***p*** | **Lower CI** | **Upper CI** |
| ***Saline*** | | | | | | |
| *A ~ W* | -15.67 | 148.12 | -15.97 | < 0.0001 | -18.22 | -13.12 |
| *D ~ W* | -15.43 | 148.12 | -15.73 | < 0.0001 | -17.98 | -12.88 |
| *D ~ A* | 0.24 | 148.12 | 0.25 | 0.99 | -2.31 | 2.79 |
| *R ~ W* | -6.74 | 149.97 | -5.84 | < 0.0001 | -9.74 | -3.74 |
| *R ~ A* | 8.93 | 149.97 | 7.73 | < 0.0001 | 5.93 | 11.93 |
| *R ~ D* | 8.69 | 149.97 | 7.52 | < 0.0001 | 5.69 | 11.69 |
| ***DOI*** | | | | | | |
| *A ~ W* | -16.26 | 148.12 | -17.52 | < 0.0001 | -18.67 | -13.84 |
| *D ~ W* | 5.36 | 148.12 | 5.78 | < 0.0001 | 2.95 | 7.77 |
| *D ~ A* | 21.62 | 148.12 | 23.30 | < 0.0001 | 19.21 | 24.03 |
| *R ~ W* | 4.11 | 149.53 | 3.95 | 0.00069 | 1.41 | 6.81 |
| *R ~ A* | 20.36 | 149.53 | 19.59 | < 0.0001 | 17.66 | 23.06 |
| *R ~ D* | -1.26 | 149.53 | -1.21 | 0.62 | -3.96 | 1.44 |
| ***DOI + Vol*** | | | | | | |
| *A ~ W* | -14.57 | 148.12 | -12.48 | < 0.0001 | -17.60 | -11.53 |
| *D ~ W* | -14.85 | 148.12 | -12.72 | < 0.0001 | -17.88 | -11.81 |
| *D ~ A* | -0.28 | 148.12 | -0.24 | 1.00 | -3.31 | 2.75 |
| *R ~ W* | -3.20 | 148.12 | -2.74 | 0.035 | -6.23 | -0.16 |
| *R ~ A* | 11.37 | 148.12 | 9.74 | < 0.0001 | 8.34 | 14.40 |
| *R ~ D* | 11.65 | 148.12 | 9.98 | < 0.0001 | 8.62 | 14.68 |

Statistical outputs include the sex effect for the linear mixed model (LMM), the results of the likelihood ratio test, and the results for each pairwise comparison between states of wake (W), anesthesia (A), drug (D), and recovery (R) for saline, DOI, or DOI + volinanserin (DOI + Vol). Corrected p values using Tukey’s method are reported for pairwise comparisons. CI = 95% confidence interval; df = degrees of freedom; Estimate = difference in means.

**Table S21 Statistical Outputs for Theta Average Node Degree before and after Infusion of Saline, DOI, or DOI with Volinanserin Pretreatment during Isoflurane Anesthesia**

| | Estimate | df | *t* | *p* | Lower CI | Upper CI |
|---|---|---|---|---|---|---|
| ***LMM: Female v. Male*** | -0.084 | 11.17 | -0.065 | 0.95 | -2.61 | 2.37 |
| | | **df** | $X^2$ | ***p*** | | |
| ***Likelihood Ratio Test*** | - | 6.00 | 46.06 | < 0.0001 | - | - |
| | **Estimate** | **df** | ***t*** | ***p*** | **Lower CI** | **Upper CI** |
| ***Saline*** | | | | | | |
| *A ~ W* | -3.74 | 113.23 | -2.94 | 0.021 | -7.07 | -0.42 |
| *D ~ W* | -4.71 | 113.23 | -3.70 | 0.0019 | -8.04 | -1.39 |
| *D ~ A* | -0.97 | 113.23 | -0.76 | 0.87 | -4.29 | 2.36 |
| *R ~ W* | -7.01 | 113.40 | -5.35 | < 0.0001 | -10.42 | -3.59 |
| *R ~ A* | -3.26 | 113.40 | -2.49 | 0.066 | -6.68 | 0.15 |
| *R ~ D* | -2.30 | 113.40 | -1.75 | 0.30 | -5.71 | 1.12 |
| ***DOI*** | | | | | | |
| *A ~ W* | -4.54 | 113.23 | -4.016 | 0.00061 | -7.49 | -1.59 |
| *D ~ W* | -14.14 | 113.23 | -12.51 | < 0.0001 | -17.09 | -11.19 |
| *D ~ A* | -9.60 | 113.23 | -8.49 | < 0.0001 | -12.55 | -6.65 |
| *R ~ W* | -7.73 | 115.63 | -6.32 | < 0.0001 | -10.91 | -4.54 |
| *R ~ A* | -3.19 | 115.63 | -2.61 | 0.050 | -6.37 | -0.0026 |
| *R ~ D* | 6.41 | 115.63 | 5.25 | < 0.0001 | 3.23 | 9.60 |
| ***DOI + Vol*** | | | | | | |
| *A ~ W* | -6.72 | 113.23 | -5.27 | < 0.0001 | -10.04 | -3.39 |
| *D ~ W* | -7.79 | 113.23 | -6.11 | < 0.0001 | -11.11 | -4.46 |
| *D ~ A* | -1.07 | 113.23 | -0.84 | 0.84 | -4.40 | 2.25 |
| *R ~ W* | -6.96 | 113.53 | -5.15 | < 0.0001 | -10.48 | -3.44 |
| *R ~ A* | -0.24 | 113.53 | -0.18 | 1.00 | -3.76 | 3.28 |
| *R ~ D* | 0.83 | 113.53 | 0.62 | 0.93 | -2.69 | 4.35 |

Statistical outputs include the sex effect for the linear mixed model (LMM), the results of the likelihood ratio test, and the results for each pairwise comparison between states of wake (W), anesthesia (A), drug (D), and recovery (R) for saline, DOI, or DOI + volinanserin (DOI + Vol). Corrected p values using Tukey's method are reported for pairwise comparisons. CI = 95% confidence interval; df = degrees of freedom; Estimate = difference in means.

**Table S22 Statistical Outputs for High Gamma Average Node Degree before and after Infusion of Saline, DOI, or DOI with Volinanserin Pretreatment during Isoflurane Anesthesia**

| | Estimate | df | *t* | *p* | Lower CI | Upper CI |
|---|---|---|---|---|---|---|
| ***LMM: Female v. Male*** | 0.76 | 11.18 | 0.62 | 0.55 | -1.68 | 3.10 |
| | | **df** | $X^2$ | ***p*** | | |
| ***Likelihood Ratio Test*** | - | 6.00 | 101.66 | < 0.0001 | - | - |
| | **Estimate** | **df** | ***t*** | ***p*** | **Lower CI** | **Upper CI** |
| ***Saline*** | | | | | | |
| *A ~ W* | -16.83 | 113.58 | -9.76 | < 0.0001 | -21.33 | -12.34 |
| *D ~ W* | -16.09 | 113.58 | -9.33 | < 0.0001 | -20.59 | -11.59 |
| *D ~ A* | 0.75 | 113.58 | 0.43 | 0.97 | -3.75 | 5.25 |
| *R ~ W* | -1.71 | 113.93 | -0.97 | 0.77 | -6.33 | 2.91 |
| *R ~ A* | 15.12 | 113.93 | 8.54 | < 0.0001 | 10.50 | 19.74 |
| *R ~ D* | 14.37 | 113.93 | 8.12 | < 0.0001 | 9.76 | 18.99 |
| ***DOI*** | | | | | | |
| *A ~ W* | -15.26 | 113.58 | -9.98 | < 0.0001 | -19.24 | -11.27 |
| *D ~ W* | 4.65 | 113.58 | 3.043 | 0.015 | 0.67 | 8.64 |
| *D ~ A* | 19.91 | 113.58 | 13.02 | < 0.0001 | 15.92 | 23.90 |
| *R ~ W* | 4.07 | 116.59 | 2.47 | 0.070 | -0.22 | 8.36 |
| *R ~ A* | 19.32 | 116.59 | 11.74 | < 0.0001 | 15.03 | 23.61 |
| *R ~ D* | -0.59 | 116.59 | -0.36 | 0.98 | -4.88 | 3.70 |
| ***DOI + Vol*** | | | | | | |
| *A ~ W* | -17.06 | 113.58 | -9.89 | < 0.0001 | -21.56 | -12.56 |
| *D ~ W* | -16.61 | 113.58 | -9.63 | < 0.0001 | -21.11 | -12.11 |
| *D ~ A* | 0.45 | 113.58 | 0.26 | 0.99 | -4.05 | 4.95 |
| *R ~ W* | -2.71 | 114.20 | -1.48 | 0.45 | -7.47 | 2.055 |
| *R ~ A* | 14.35 | 114.20 | 7.86 | < 0.0001 | 9.59 | 19.11 |
| *R ~ D* | 13.91 | 114.20 | 7.62 | < 0.0001 | 9.15 | 18.67 |

Statistical outputs include the sex effect for the linear mixed model (LMM), the results of the likelihood ratio test, and the results for each pairwise comparison between states of wake (W), anesthesia (A), drug (D), and recovery (R) for saline, DOI, or DOI + volinanserin (DOI + Vol). Corrected p values using Tukey's method are reported for pairwise comparisons. CI = 95% confidence interval; df = degrees of freedom; Estimate = difference in means.

**Table S23 Statistical Outputs for Active and Passive Emergence after the Infusion of Saline, DOI, or Lisuride during Propofol Anesthesia**

| ***Arousal Score*** | **Estimate** | **df** | ***t*** | ***p*** | **Lower CI** | **Upper CI** |
|---|---|---|---|---|---|---|
| *LMM: Female v. Male* | -0.25 | 28.00 | -2.02 | 0.053 | -0.49 | -0.017 |
| *DOI ~ Saline* | 2.99 | 18.94 | 18.33 | < 0.0001 | 2.57 | 3.40 |
| *Lisuride ~ Saline* | -0.035 | 20.17 | -0.23 | 0.97 | -0.42 | 0.35 |
| *Lisuride ~ DOI* | -3.024 | 20.91 | -19.47 | < 0.0001 | -3.42 | -2.63 |
| ***RORR*** | **Estimate** | **df** | ***t*** | ***p*** | **Lower CI** | **Upper CI** |
| *LMM: Female v. Male* | 251.21 | 15.91 | 2.73 | 0.015 | 76.64 | 424.62 |
| *DOI ~ Saline* | -126.77 | 15.48 | -1.01 | 0.58 | -452.91 | 199.37 |
| *Lisuride ~ Saline* | 217.68 | 15.31 | 2.35 | 0.078 | -21.987 | 457.33 |
| *Lisuride ~ DOI* | 344.44 | 14.65 | 2.83 | 0.033 | 27.16 | 661.73 |

Statistical outputs include the sex effect for the linear mixed model (LMM) and results for each pairwise comparison between conditions (saline, DOI, and lisuride) for arousal score (i.e., active emergence) and time to the return of righting reflex (RORR) after cessation of the anesthetic (i.e., passive emergence). Corrected p values using Tukey's method are reported for pairwise comparisons. CI = 95% confidence interval; df = degrees of freedom; Estimate = difference in means.

**Table S24 Statistical Outputs for Theta Relative Power before and after Infusion of Saline, DOI, or Lisuride during Propofol Anesthesia**

| | Estimate | df | *t* | *p* | Lower CI | Upper CI |
|---|---|---|---|---|---|---|
| ***LMM: Female v. Male*** | 0.16 | 7.33 | 1.21 | 0.27 | -0.13 | 0.41 |
| | | **df** | $X^2$ | ***p*** | | |
| ***Likelihood Ratio Test*** | - | 6.00 | 23.84 | 0.00056 | - | - |
| | **Estimate** | **df** | ***t*** | ***p*** | **Lower CI** | **Upper CI** |
| ***Saline*** | | | | | | |
| *A ~ W* | -0.85 | 94.73 | -3.18 | 0.010 | -1.55 | -0.15 |
| *D ~ W* | -0.77 | 94.73 | -2.89 | 0.024 | -1.47 | -0.073 |
| *D ~ A* | 0.078 | 94.73 | 0.29 | 0.99 | -0.62 | 0.78 |
| *R ~ W* | -1.37 | 94.73 | -5.16 | < 0.0001 | -2.07 | -0.68 |
| *R ~ A* | -0.53 | 94.73 | -1.97 | 0.21 | -1.22 | 0.17 |
| *R ~ D* | -0.60 | 94.73 | -2.27 | 0.11 | -1.30 | 0.093 |
| ***DOI*** | | | | | | |
| *A ~ W* | -1.011 | 94.73 | -3.79 | 0.0015 | -1.71 | -0.31 |
| *D ~ W* | -1.41 | 94.73 | -5.31 | < 0.0001 | -2.11 | -0.72 |
| *D ~ A* | -0.40 | 94.73 | -1.51 | 0.43 | -1.10 | 0.29 |
| *R ~ W* | -1.30 | 95.18 | -4.06 | 0.00058 | -2.13 | -0.46 |
| *R ~ A* | -0.28 | 95.18 | -0.89 | 0.81 | -1.12 | 0.55 |
| *R ~ D* | 0.12 | 95.18 | 0.37 | 0.98 | -0.72 | 0.95 |
| ***Lisuride*** | | | | | | |
| *A ~ W* | -0.99 | 94.73 | -4.45 | 0.00014 | -1.57 | -0.41 |
| *D ~ W* | -0.28 | 94.73 | -1.28 | 0.58 | -0.86 | 0.30 |
| *D ~ A* | 0.70 | 94.73 | 3.17 | 0.011 | 0.12 | 1.28 |
| *R ~ W* | -1.90 | 95.70 | -8.20 | < 0.0001 | -2.51 | -1.30 |
| *R ~ A* | -0.92 | 95.70 | -3.95 | 0.00083 | -1.53 | -0.31 |
| *R ~ D* | -1.62 | 95.70 | -6.98 | < 0.0001 | -2.23 | -1.01 |

Statistical outputs include the sex effect for the linear mixed model (LMM), the results of the likelihood ratio test, and the results for each pairwise comparison between states of wake (W), anesthesia (A), drug (D), and recovery (R) for saline, DOI, or lisuride. Corrected p values using Tukey's method are reported for pairwise comparisons. CI = 95% confidence interval; df = degrees of freedom; Estimate = difference in means.

**Table S25 Statistical Outputs for High Gamma Relative Power before and after Infusion of Saline, DOI, or Lisuride during Propofol Anesthesia**

| | Estimate | df | *t* | *p* | Lower CI | Upper CI |
|---|---|---|---|---|---|---|
| ***LMM: Female v. Male*** | -0.00036 | 11.71 | -0.61 | 0.55 | -0.0015 | 0.00076 |
| | | **df** | $X^2$ | *p* | | |
| ***Likelihood Ratio Test*** | - | 6.00 | 25.070 | 0.00033 | - | - |
| | **Estimate** | **df** | ***t*** | ***p*** | **Lower CI** | **Upper CI** |
| ***Saline*** | | | | | | |
| *A ~ W* | -0.0074 | 93.82 | -6.28 | < 0.0001 | -0.011 | -0.0043 |
| *D ~ W* | -0.0074 | 93.82 | -6.30 | < 0.0001 | -0.011 | -0.0043 |
| *D ~ A* | 0.000017 | 93.82 | -0.014 | 1.00 | -0.0031 | 0.0031 |
| *R ~ W* | -0.0029 | 93.82 | -2.47 | 0.072 | -0.0060 | 0.00017 |
| *R ~ A* | 0.0045 | 93.82 | 3.82 | 0.0014 | 0.0014 | 0.0076 |
| *R ~ D* | 0.0045 | 93.82 | 3.83 | 0.0013 | 0.0014 | 0.0076 |
| ***DOI*** | | | | | | |
| *A ~ W* | -0.0076 | 93.82 | -6.42 | < 0.0001 | -0.011 | -0.0045 |
| *D ~ W* | -0.0043 | 93.82 | -3.66 | 0.0023 | -0.0074 | -0.0012 |
| *D ~ A* | 0.0033 | 93.82 | 2.77 | 0.034 | 0.00018 | 0.0063 |
| *R ~ W* | 0.0036 | 94.25 | 2.52 | 0.063 | -0.00013 | 0.0072 |
| *R ~ A* | 0.011 | 94.25 | 7.89 | < 0.0001 | 0.0074 | 0.015 |
| *R ~ D* | 0.0079 | 94.25 | 5.58 | < 0.0001 | 0.0042 | 0.012 |
| ***Lisuride*** | | | | | | |
| *A ~ W* | -0.0086 | 94.73 | -8.60 | < 0.0001 | -0.011 | -0.0060 |
| *D ~ W* | -0.0085 | 94.73 | -8.49 | < 0.0001 | -0.011 | -0.0059 |
| *D ~ A* | 0.00010 | 93.82 | 0.11 | 1.00 | -0.0025 | 0.0027 |
| *R ~ W* | -0.0039 | 95.78 | -3.75 | 0.0017 | -0.0067 | -0.0012 |
| *R ~ A* | 0.0047 | 94.78 | 4.56 | < 0.0001 | 0.0020 | 0.0074 |
| *R ~ D* | 0.0046 | 94.78 | 4.46 | 0.00013 | 0.0019 | 0.0073 |

Statistical outputs include the sex effect for the linear mixed model (LMM), the results of the likelihood ratio test, and the results for each pairwise comparison between states of wake (W), anesthesia (A), drug (D), and recovery (R) for saline, DOI, or lisuride. Corrected p values using Tukey's method are reported for pairwise comparisons. CI = 95% confidence interval; df = degrees of freedom; Estimate = difference in means.

**Table S26 Statistical Outputs for Theta Global Coherence before and after Infusion of Saline, DOI, or Lisuride during Propofol Anesthesia**

| | **Estimate** | **df** | ***t*** | ***p*** | **Lower CI** | **Upper CI** |
|---|---|---|---|---|---|---|
| ***LMM: Female v. Male*** | -0.063 | 10.55 | -0.59 | 0.57 | -0.27 | 0.15 |
| | | **df** | ***$X^2$*** | ***p*** | | |
| ***Likelihood Ratio Test*** | - | 6.00 | 19.61 | 0.011 | - | - |
| | **Estimate** | **df** | ***t*** | ***p*** | **Lower CI** | **Upper CI** |
| ***Saline*** | | | | | | |
| *A ~ W* | -1.02 | 84.35 | -5.93 | < 0.0001 | -1.46 | -0.57 |
| *D ~ W* | -1.07 | 84.35 | -6.26 | < 0.0001 | -1.52 | -0.62 |
| *D ~ A* | -0.057 | 84.35 | -0.33 | 0.99 | -0.51 | 0.39 |
| *R ~ W* | -0.80 | 84.35 | -4.68 | < 0.0001 | -1.25 | -0.35 |
| *R ~ A* | 0.21 | 84.35 | 1.25 | 0.60 | -0.24 | 0.66 |
| *R ~ D* | 0.27 | 84.35 | 1.58 | 0.40 | -0.18 | 0.72 |
| ***DOI*** | | | | | | |
| *A ~ W* | -1.07 | 84.35 | -6.26 | < 0.0001 | -1.52 | -0.62 |
| *D ~ W* | -1.77 | 84.35 | -10.31 | < 0.0001 | -2.22 | -1.32 |
| *D ~ A* | -0.70 | 84.35 | -4.06 | 0.00063 | -1.14 | -0.25 |
| *R ~ W* | -0.95 | 84.67 | -4.79 | < 0.0001 | -1.46 | -0.43 |
| *R ~ A* | 0.13 | 84.67 | 0.64 | 0.92 | -0.39 | 0.64 |
| *R ~ D* | 0.82 | 84.67 | 4.16 | 0.00043 | 0.30 | 1.34 |
| ***Lisuride*** | | | | | | |
| *A ~ W* | -0.99 | 84.35 | -7.06 | < 0.0001 | -1.35 | -0.62 |
| *D ~ W* | -1.04 | 84.35 | -7.44 | < 0.0001 | -1.41 | -0.67 |
| *D ~ A* | -0.054 | 84.35 | -0.38 | 0.98 | -0.42 | 0.31 |
| *R ~ W* | -0.89 | 85.02 | -6.04 | < 0.0001 | -1.28 | -0.50 |
| *R ~ A* | 0.098 | 85.02 | 0.66 | 0.91 | -0.29 | 0.48 |
| *R ~ D* | 0.15 | 85.02 | 1.03 | 0.73 | -0.23 | 0.54 |

Statistical outputs include the sex effect for the linear mixed model (LMM), the results of the likelihood ratio test, and the results for each pairwise comparison between states of wake (W), anesthesia (A), drug (D), and recovery (R) for saline, DOI, or lisuride. Corrected p values using Tukey's method are reported for pairwise comparisons. CI = 95% confidence interval; df = degrees of freedom; Estimate = difference in means.

**Table S27 Statistical Outputs for High Gamma Global Coherence before and after Infusion of Saline, DOI, or Lisuride during Propofol Anesthesia**

| | Estimate | df | *t* | *p* | Lower CI | Upper CI |
|---|---|---|---|---|---|---|
| ***LMM: Female v. Male*** | 0.059 | 11.47 | 0.66 | 0.52 | -0.11 | 0.23 |
| | | **df** | ***X*²** | ***p*** | | |
| ***Likelihood Ratio Test*** | - | 6.00 | 111.01 | < 0.0001 | - | - |
| | **Estimate** | **df** | ***t*** | ***p*** | **Lower CI** | **Upper CI** |
| ***Saline*** | | | | | | |
| *A ~ W* | -1.34 | 84.51 | -8.44 | < 0.0001 | -1.76 | -0.93 |
| *D ~ W* | -1.33 | 84.51 | -8.32 | < 0.0001 | -1.74 | -0.91 |
| *D ~ A* | 0.019 | 84.51 | 0.12 | 1.00 | -0.40 | 0.44 |
| *R ~ W* | -0.40 | 84.51 | -2.48 | 0.070 | -0.81 | 0.022 |
| *R ~ A* | 0.95 | 84.51 | 5.96 | < 0.0001 | 0.53 | 1.37 |
| *R ~ D* | 0.93 | 84.51 | 5.84 | < 0.0001 | 0.51 | 1.35 |
| ***DOI*** | | | | | | |
| *A ~ W* | -1.46 | 84.51 | -9.15 | < 0.0001 | -1.87 | -1.04 |
| *D ~ W* | 0.50 | 84.51 | 3.13 | 0.013 | 0.081 | 0.92 |
| *D ~ A* | 1.95 | 84.51 | 12.27 | < 0.0001 | 1.54 | 2.37 |
| *R ~ W* | 0.30 | 84.87 | 1.61 | 0.38 | -0.19 | 0.78 |
| *R ~ A* | 1.75 | 84.87 | 9.55 | < 0.0001 | 1.27 | 2.23 |
| *R ~ D* | -0.20 | 84.87 | -1.10 | 0.69 | -0.68 | 0.28 |
| ***Lisuride*** | | | | | | |
| *A ~ W* | -1.45 | 84.51 | -11.17 | < 0.0001 | -1.79 | -1.11 |
| *D ~ W* | -1.83 | 84.51 | -14.07 | < 0.0001 | -2.17 | -1.49 |
| *D ~ A* | -0.38 | 84.51 | -2.90 | 0.024 | -0.72 | -0.037 |
| *R ~ W* | -0.66 | 85.34 | -4.80 | < 0.0001 | -1.016 | -0.30 |
| *R ~ A* | 0.80 | 85.34 | 5.81 | < 0.0001 | 0.44 | 1.15 |
| *R ~ D* | 1.17 | 85.34 | 8.56 | < 0.0001 | 0.81 | 1.53 |

Statistical outputs include the sex effect for the linear mixed model (LMM), the results of the likelihood ratio test, and the results for each pairwise comparison between states of wake (W), anesthesia (A), drug (D), and recovery (R) for saline, DOI, or lisuride. Corrected p values using Tukey's method are reported for pairwise comparisons. CI = 95% confidence interval; df = degrees of freedom; Estimate = difference in means.

**Table S28 Statistical Outputs for Theta Feedforward Connectivity before and after Infusion of Saline, DOI, or Lisuride during Propofol Anesthesia**

| | Estimate | df | *t* | *p* | Lower CI | Upper CI |
|---|---|---|---|---|---|---|
| ***LMM: Female v. Male*** | -0.0020 | 11.29 | -0.65 | 0.53 | -0.0078 | 0.0038 |
| | | **df** | **$X^2$** | ***p*** | | |
| ***Likelihood Ratio Test*** | - | 6.00 | 20.46 | 0.0023 | - | - |
| | **Estimate** | **df** | ***t*** | ***p*** | **Lower CI** | **Upper CI** |
| ***Saline*** | | | | | | |
| *A ~ W* | -0.025 | 84.63 | -4.34 | 0.00023 | -0.041 | -0.010 |
| *D ~ W* | -0.027 | 84.63 | -4.54 | 0.00011 | -0.042 | -0.011 |
| *D ~ A* | -0.0012 | 84.63 | -0.20 | 1.00 | -0.017 | 0.014 |
| *R ~ W* | -0.021 | 84.63 | -3.51 | 0.0039 | -0.036 | -0.0052 |
| *R ~ A* | 0.0048 | 84.63 | 0.82 | 0.84 | -0.011 | 0.020 |
| *R ~ D* | 0.0060 | 84.63 | 1.02 | 0.74 | -0.0094 | 0.021 |
| ***DOI*** | | | | | | |
| *A ~ W* | -0.029 | 84.63 | -5.00 | < 0.0001 | -0.045 | -0.014 |
| *D ~ W* | -0.046 | 84.63 | -7.77 | < 0.0001 | -0.061 | -0.030 |
| *D ~ A* | -0.016 | 84.63 | -2.77 | 0.034 | -0.032 | -0.00087 |
| *R ~ W* | -0.012 | 85.01 | -1.75 | 0.31 | -0.029 | 0.0059 |
| *R ~ A* | 0.018 | 85.01 | 2.60 | 0.053 | 0.00015 | 0.035 |
| *R ~ D* | 0.034 | 85.01 | 5.00 | < 0.0001 | 0.016 | 0.051 |
| ***Lisuride*** | | | | | | |
| *A ~ W* | -0.030 | 84.63 | -6.18 | < 0.0001 | -0.042 | -0.017 |
| *D ~ W* | -0.026 | 84.63 | -5.32 | < 0.0001 | -0.038 | -0.013 |
| *D ~ A* | 0.0041 | 84.63 | 0.85 | 0.83 | -0.0085 | 0.017 |
| *R ~ W* | -0.028 | 85.55 | -5.45 | < 0.0001 | -0.041 | -0.014 |
| *R ~ A* | 0.0021 | 85.55 | 0.42 | 0.98 | -0.011 | 0.015 |
| *R ~ D* | -0.0020 | 85.55 | -0.39 | 0.98 | -0.015 | 0.011 |

Statistical outputs include the sex effect for the linear mixed model (LMM), the results of the likelihood ratio test, and the results for each pairwise comparison between states of wake (W), anesthesia (A), drug (D), and recovery (R) for saline, DOI, or lisuride. Corrected p values using Tukey's method are reported for pairwise comparisons. CI = 95% confidence interval; df = degrees of freedom; Estimate = difference in means.

**Table S29 Statistical Outputs for High Gamma Feedforward Connectivity before and after Infusion of Saline, DOI, or Lisuride during Propofol Anesthesia**

| | **Estimate** | **df** | ***t*** | ***p*** | **Lower CI** | **Upper CI** |
|---|---|---|---|---|---|---|
| ***LMM: Female v. Male*** | 0.00081 | 10.54 | 0.97 | 0.51 | -0.0016 | 0.0031 |
| | | **df** | $X^2$ | ***p*** | | |
| ***Likelihood Ratio Test*** | - | 6.00 | 56.61 | < 0.0001 | - | - |
| | **Estimate** | **df** | ***t*** | ***p*** | **Lower CI** | **Upper CI** |
| ***Saline*** | | | | | | |
| *A ~ W* | -0.020 | 84.45 | -9.53 | < 0.0001 | -0.025 | -0.014 |
| *D ~ W* | -0.020 | 84.45 | -9.37 | < 0.0001 | -0.025 | -0.014 |
| *D ~ A* | 0.00034 | 84.45 | 0.16 | 1.00 | -0.0051 | 0.0058 |
| *R ~ W* | -0.013 | 84.45 | -6.11 | < 0.0001 | -0.018 | -0.0073 |
| *R ~ A* | 0.0071 | 84.45 | 3.42 | 0.0053 | 0.0017 | 0.013 |
| *R ~ D* | 0.0068 | 84.45 | 3.26 | 0.0086 | 0.0013 | 0.012 |
| ***DOI*** | | | | | | |
| *A ~ W* | -0.020 | 84.45 | -9.55 | < 0.0001 | -0.025 | -0.014 |
| *D ~ W* | -0.0034 | 84.45 | -1.63 | 0.37 | -0.0089 | 0.0021 |
| *D ~ A* | 0.017 | 84.45 | 7.92 | < 0.0001 | 0.011 | 0.022 |
| *R ~ W* | -0.0081 | 84.80 | -3.37 | 0.0061 | -0.014 | -0.0018 |
| *R ~ A* | 0.012 | 84.80 | 4.92 | < 0.0001 | 0.0055 | 0.018 |
| *R ~ D* | -0.0047 | 84.80 | -1.95 | 0.22 | -0.011 | 0.0016 |
| ***Lisuride*** | | | | | | |
| *A ~ W* | -0.017 | 84.45 | -10.02 | < 0.0001 | -0.021 | -0.013 |
| *D ~ W* | -0.019 | 84.45 | -11.38 | < 0.0001 | -0.024 | -0.015 |
| *D ~ A* | -0.0023 | 84.45 | -1.36 | 0.53 | -0.0068 | 0.0021 |
| *R ~ W* | -0.0082 | 85.22 | -4.59 | < 0.0001 | -0.013 | -0.0035 |
| *R ~ A* | 0.0088 | 85.22 | 4.92 | < 0.0001 | 0.0041 | 0.013 |
| *R ~ D* | 0.011 | 85.22 | 6.21 | < 0.0001 | 0.0064 | 0.016 |

Statistical outputs include the sex effect for the linear mixed model (LMM), the results of the likelihood ratio test, and the results for each pairwise comparison between states of wake (W), anesthesia (A), drug (D), and recovery (R) for saline, DOI, or lisuride. Corrected p values using Tukey's method are reported for pairwise comparisons. CI = 95% confidence interval; df = degrees of freedom; Estimate = difference in means.

**Table S30 Statistical Outputs for Theta Feedback Connectivity before and after Infusion of Saline, DOI, or Lisuride during Propofol Anesthesia**

| | Estimate | df | *t* | *p* | Lower CI | Upper CI |
|---|---|---|---|---|---|---|
| ***LMM: Female v. Male*** | -0.0017 | 94.00 | -0.70 | 0.48 | -0.0061 | 0.0027 |
| | | **df** | $X^2$ | ***p*** | | |
| ***Likelihood Ratio Test*** | - | 6.00 | 5.80 | 0.45 | - | - |
| | **Estimate** | **df** | ***t*** | ***p*** | **Lower CI** | **Upper CI** |
| ***Saline*** | | | | | | |
| *A ~ W* | -0.026 | 85.20 | -4.34 | 0.00022 | -0.041 | -0.010 |
| *D ~ W* | -0.026 | 85.20 | -4.35 | 0.00021 | -0.041 | -0.010 |
| *D ~ A* | 0.000066 | 85.20 | -0.011 | 1.00 | -0.016 | 0.015 |
| *R ~ W* | -0.016 | 85.20 | -2.74 | 0.037 | -0.032 | -0.00070 |
| *R ~ A* | 0.0095 | 85.20 | 1.60 | 0.38 | -0.0060 | 0.025 |
| *R ~ D* | 0.010 | 85.20 | 1.62 | 0.38 | -0.0059 | 0.025 |
| ***DOI*** | | | | | | |
| *A ~ W* | -0.025 | 85.20 | -4.31 | 0.00026 | -0.041 | -0.010 |
| *D ~ W* | -0.034 | 85.20 | -5.72 | < 0.0001 | -0.049 | -0.018 |
| *D ~ A* | -0.0083 | 85.20 | -1.41 | 0.49 | -0.024 | 0.0071 |
| *R ~ W* | -0.011 | 85.62 | -1.59 | 0.39 | -0.029 | 0.0070 |
| *R ~ A* | 0.015 | 85.62 | 2.15 | 0.15 | -0.0032 | 0.032 |
| *R ~ D* | 0.023 | 85.62 | 3.38 | 0.0059 | 0.0051 | 0.041 |
| ***Lisuride*** | | | | | | |
| *A ~ W* | -0.022 | 85.20 | -4.54 | 0.00011 | -0.035 | -0.0093 |
| *D ~ W* | -0.019 | 85.20 | -4.00 | 0.00078 | -0.032 | -0.0066 |
| *D ~ A* | 0.0026 | 85.20 | 0.55 | 0.95 | -0.010 | 0.015 |
| *R ~ W* | -0.011 | 86.57 | -2.20 | 0.13 | -0.024 | 0.0021 |
| *R ~ A* | 0.011 | 86.57 | 2.12 | 0.16 | -0.0025 | 0.024 |
| *R ~ D* | 0.0081 | 86.57 | 1.60 | 0.39 | -0.0052 | 0.021 |

Statistical outputs include the sex effect for the linear mixed model (LMM), the results of the likelihood ratio test, and the results for each pairwise comparison between states of wake (W), anesthesia (A), drug (D), and recovery (R) for saline, DOI, or lisuride. Corrected p values using Tukey's method are reported for pairwise comparisons. CI = 95% confidence interval; df = degrees of freedom; Estimate = difference in means.

**Table S31 Statistical Outputs for High Gamma Feedback Connectivity before and after Infusion of Saline, DOI, or Lisuride during Propofol Anesthesia**

| | Estimate | df | *t* | *p* | Lower CI | Upper CI |
|---|---|---|---|---|---|---|
| ***LMM: Female v. Male*** | 0.00081 | 8.49 | 0.53 | 0.61 | -0.0023 | 0.0036 |
| | | **df** | **$X^2$** | ***p*** | | |
| ***Likelihood Ratio Test*** | - | 6.00 | 42.062 | < 0.0001 | - | - |
| | **Estimate** | **df** | ***t*** | ***p*** | **Lower CI** | **Upper CI** |
| ***Saline*** | | | | | | |
| *A ~ W* | -0.019 | 84.28 | -8.38 | < 0.0001 | -0.025 | -0.013 |
| *D ~ W* | -0.019 | 84.28 | -8.46 | < 0.0001 | -0.025 | -0.013 |
| *D ~ A* | -0.00019 | 84.28 | -0.085 | 1.00 | -0.0062 | 0.0058 |
| *R ~ W* | -0.014 | 84.28 | -6.01 | < 0.0001 | -0.020 | -0.0077 |
| *R ~ A* | 0.0054 | 84.28 | 2.37 | 0.092 | -0.00059 | 0.011 |
| *R ~ D* | 0.0056 | 84.28 | 2.45 | 0.075 | 0.00039 | 0.012 |
| ***DOI*** | | | | | | |
| *A ~ W* | -0.019 | 84.28 | -8.55 | < 0.0001 | -0.025 | -0.013 |
| *D ~ W* | -0.0051 | 84.28 | -2.25 | 0.12 | -0.011 | 0.00084 |
| *D ~ A* | 0.014 | 84.28 | 6.30 | < 0.0001 | 0.0084 | 0.020 |
| *R ~ W* | -0.012 | 84.57 | -4.47 | 0.00014 | -0.019 | -0.0048 |
| *R ~ A* | 0.0077 | 84.57 | 2.95 | 0.021 | 0.00088 | 0.015 |
| *R ~ D* | -0.0066 | 84.57 | -2.51 | 0.065 | -0.013 | 0.00028 |
| ***Lisuride*** | | | | | | |
| *A ~ W* | -0.017 | 84.28 | -9.18 | < 0.0001 | -0.022 | -0.012 |
| *D ~ W* | -0.019 | 84.28 | -10.07 | < 0.0001 | -0.024 | -0.014 |
| *D ~ A* | -0.0016 | 84.28 | -0.88 | 0.82 | -0.0065 | 0.0032 |
| *R ~ W* | -0.0084 | 84.87 | -4.27 | 0.00029 | -0.013 | -0.0032 |
| *R ~ A* | 0.0087 | 84.87 | 4.44 | 0.00015 | 0.0036 | 0.014 |
| *R ~ D* | 0.010 | 84.87 | 5.28 | < 0.0001 | 0.0052 | 0.015 |

Statistical outputs include the sex effect for the linear mixed model (LMM), the results of the likelihood ratio test, and the results for each pairwise comparison between states of wake (W), anesthesia (A), drug (D), and recovery (R) for saline, DOI, or lisuride. Corrected p values using Tukey's method are reported for pairwise comparisons. CI = 95% confidence interval; df = degrees of freedom; Estimate = difference in means.

**Table S32 Statistical Outputs for Theta Average Node Degree before and after Infusion of Saline, DOI, or Lisuride during Propofol Anesthesia**

| | Estimate | df | *t* | *p* | Lower CI | Upper CI |
|---|---|---|---|---|---|---|
| ***LMM: Female v. Male*** | -0.41 | 11.22 | -0.45 | 0.66 | -2.18 | 1.32 |
| | | **df** | $X^2$ | ***p*** | | |
| ***Likelihood Ratio Test*** | - | 6.00 | 13.21 | 0.040 | - | - |
| | **Estimate** | **df** | ***t*** | ***p*** | **Lower CI** | **Upper CI** |
| ***Saline*** | | | | | | |
| *A ~ W* | -7.06 | 84.59 | -4.10 | 0.00055 | -11.58 | -2.54 |
| *D ~ W* | -8.05 | 84.59 | -4.67 | < 0.0001 | -12.57 | -3.53 |
| *D ~ A* | -0.99 | 84.59 | -0.57 | 0.94 | -5.51 | 3.53 |
| *R ~ W* | -5.25 | 84.59 | -3.04 | 0.016 | -9.77 | -0.73 |
| *R ~ A* | 1.81 | 84.59 | 1.05 | 0.72 | -2.70 | 6.33 |
| *R ~ D* | 2.80 | 84.59 | 1.62 | 0.37 | -1.72 | 7.32 |
| ***DOI*** | | | | | | |
| *A ~ W* | -7.79 | 84.59 | -4.52 | 0.00012 | -12.31 | -3.27 |
| *D ~ W* | -14.66 | 84.59 | -8.50 | < 0.0001 | -19.18 | -10.14 |
| *D ~ A* | -6.87 | 84.59 | -3.98 | 0.00081 | -11.39 | -2.35 |
| *R ~ W* | -6.81 | 84.96 | -3.43 | 0.0051 | -12.01 | -1.61 |
| *R ~ A* | 0.99 | 84.96 | 0.50 | 0.96 | -4.22 | 6.19 |
| *R ~ D* | 7.85 | 84.96 | 3.96 | 0.00089 | 2.65 | 13.06 |
| ***Lisuride*** | | | | | | |
| *A ~ W* | -8.18 | 84.59 | -5.81 | < 0.0001 | -11.87 | -4.49 |
| *D ~ W* | -8.95 | 84.59 | -6.36 | < 0.0001 | -12.64 | -5.26 |
| *D ~ A* | -0.77 | 84.59 | -0.55 | 0.95 | -4.46 | 2.92 |
| *R ~ W* | -6.60 | 85.47 | -4.45 | 0.00015 | -10.48 | -2.71 |
| *R ~ A* | 1.58 | 85.47 | 1.07 | 0.71 | -2.31 | 5.46 |
| *R ~ D* | 2.35 | 85.47 | 1.58 | 0.39 | -1.53 | 6.23 |

Statistical outputs include the sex effect for the linear mixed model (LMM), the results of the likelihood ratio test, and the results for each pairwise comparison between states of wake (W), anesthesia (A), drug (D), and recovery (R) for saline, DOI, or lisuride. Corrected p values using Tukey’s method are reported for pairwise comparisons. CI = 95% confidence interval; df = degrees of freedom; Estimate = difference in means.

**Table S33 Statistical Outputs for High Gamma Average Node Degree before and after Infusion of Saline, DOI, or Lisuride during Propofol Anesthesia**

| | Estimate | df | *t* | *p* | Lower CI | Upper CI |
|---|---|---|---|---|---|---|
| ***LMM: Female v. Male*** | 0.47 | 9.65 | 0.56 | 0.59 | -1.19 | 2.06 |
| | | **df** | $X^2$ | *p* | | |
| ***Likelihood Ratio Test*** | - | 6 | 116.30 | < 0.0001 | - | - |
| | **Estimate** | **df** | ***t*** | ***p*** | **Lower CI** | **Upper CI** |
| ***Saline*** | | | | | | |
| *A ~ W* | -16.14 | 84.46 | -11.02 | < 0.0001 | -19.97 | -12.30 |
| *D ~ W* | -16.07 | 84.46 | -10.97 | < 0.0001 | -19.91 | -12.23 |
| *D ~ A* | 0.064 | 84.46 | 0.044 | 1.00 | -3.78 | 3.90 |
| *R ~ W* | -3.25 | 84.46 | -2.22 | 0.13 | -7.09 | 0.59 |
| *R ~ A* | 12.88 | 84.46 | 8.79 | < 0.0001 | 9.04 | 16.72 |
| *R ~ D* | 12.82 | 84.46 | 8.75 | < 0.0001 | 8.98 | 16.66 |
| ***DOI*** | | | | | | |
| *A ~ W* | -16.34 | 84.46 | -11.16 | < 0.0001 | -20.18 | -12.50 |
| *D ~ W* | 3.30 | 84.46 | 2.26 | 0.12 | -0.54 | 7.14 |
| *D ~ A* | 19.65 | 84.46 | 13.41 | < 0.0001 | 15.81 | 23.49 |
| *R ~ W* | 2.55 | 84.80 | 1.51 | 0.43 | -1.87 | 6.98 |
| *R ~ A* | 18.90 | 84.80 | 11.20 | < 0.0001 | 14.48 | 23.32 |
| *R ~ D* | -0.75 | 84.80 | -0.44 | 0.97 | -5.17 | 3.67 |
| ***Lisuride*** | | | | | | |
| *A ~ W* | -16.64 | 84.46 | -13.91 | < 0.0001 | -19.77 | -13.50 |
| *D ~ W* | -18.64 | 84.46 | -15.59 | < 0.0001 | -21.78 | -15.51 |
| *D ~ A* | -2.01 | 84.46 | -1.68 | 0.34 | -5.14 | 1.13 |
| *R ~ W* | -6.29 | 85.23 | -4.99 | < 0.0001 | -9.59 | -2.99 |
| *R ~ A* | 10.35 | 85.23 | 8.21 | < 0.0001 | 7.05 | 13.65 |
| *R ~ D* | 12.35 | 85.23 | 9.81 | < 0.0001 | 9.05 | 15.66 |

Statistical outputs include the sex effect for the linear mixed model (LMM), the results of the likelihood ratio test, and the results for each pairwise comparison between states of wake (W), anesthesia (A), drug (D), and recovery (R) for saline, DOI, or lisuride. Corrected p values using Tukey's method are reported for pairwise comparisons. CI = 95% confidence interval; df = degrees of freedom; Estimate = difference in means.

**Movie S1.**

Video depicting behavioral arousal after intravenous infusion of DOI (0.5 mg/kg) during propofol anesthesia. Rat initially is unmoving during propofol anesthesia. The rat begins showing signs of behavioral arousal during propofol anesthesia by the end of the 1-minute infusion of DOI. Following a short time skip, the rat achieves return of the righting reflex, a surrogate for return of consciousness in rodents, 2 minutes and 12 seconds after the end of the DOI infusion. The return of righting reflex is when the rat, previously placed on its back in the supine position, rights itself on all four paws. Note that the rat is receiving propofol anesthesia for the duration of the video.

**Movie S2.**

Video depicting behavioral arousal after intravenous infusion of DOI (0.5 mg/kg) during isoflurane anesthesia. The rat is initially unmoving during isoflurane anesthesia. Towards the end of the 1-minute DOI infusion, the rat begins showing signs of behavioral arousal during isoflurane anesthesia and achieves return of the righting reflex, a surrogate for return of consciousness in rodents, 10 seconds after the end of the DOI infusion. The return of righting reflex is when the rat, previously placed on its back in the supine position, rights itself on all four paws. Note that the rat is receiving isoflurane anesthesia for the duration of the video.
